# Somatic transposition in the *Drosophila* intestine occurs in active chromatin and is associated with tumor suppressor gene inactivation

**DOI:** 10.1101/2020.07.10.166629

**Authors:** Katarzyna Siudeja, Marius van den Beek, Nick Riddiford, Benjamin Boumard, Annabelle Wurmser, Marine Stefanutti, Sonia Lameiras, Allison J. Bardin

**Author notes:** Equal contribution (Katarzyna Siudeja, Marius van den Beek). Authors for correspondence (,) Twitter: @BardinLab.

## Abstract

Transposable elements (TEs) play a significant role in evolution by contributing to genetic variation through germline insertional activity. However, how TEs act in somatic cells and tissues is not well understood. Here, we address the prevalence of transposition in a somatic tissue, exploiting the *Drosophila* midgut as a model system. Using whole-genome sequencing of *in vivo* clonally expanded gut tissue, we map hundreds of high-confidence somatic TE integration sites genome-wide. We show that somatic retrotransposon insertions are associated with inactivation of the tumor suppressor *Notch*, likely contributing to neoplasia formation. Moreover, by applying Oxford Nanopore long-read sequencing technology, as well as by mapping germline TE activity, we provide evidence suggesting tissue-specific differences in retrotransposition. By comparing somatic TE insertional activity with transcriptomic and small RNA sequencing data, we demonstrate that transposon mobility cannot be simply predicted by whole tissue TE expression levels or by small RNA pathway activity. Finally, we reveal that somatic TE insertions in the adult fly intestine are found preferentially in genic regions and open, transcriptionally active chromatin. Together, our findings provide clear evidence of ongoing somatic transposition in *Drosophila* and delineate previously unknown underlying features of somatic TE mobility *in vivo*.

## Introduction

Transposable elements (TEs) are DNA sequences that shape evolution through their capacity to amplify and mobilize, thereby altering the structural and regulatory landscape of the genome. Numerous mechanisms restrict the mobility of TEs and therefore, their mutagenic potential. In germline and somatic cells, TE silencing is achieved by chromatin modifications and small RNA-directed degradation of TE transcripts (Molaro and Malik 2016; Deniz et al. 2019; Cosby et al. 2019). The escape of TEs from silencing allows their propagation in the genome. While *de novo* TE insertions in the germline are relatively easy to detect as they result in heritable genomic changes that can be detected through sequencing, TE mobility in somatic cells is more difficult to study. Indeed, the heterogeneity of transposition events within somatic tissues imposes technical challenges as rare TE insertion events affecting a subpopulation of cells often fall below the limits of detection. Thus, the degree to which TEs evade silencing and contribute to somatic genome alteration is much less well understood in developing and adult tissues.

Nonetheless, evidence for active somatic transposition has been recently mounting. Reporters of transposon activity suggested TE mobility in neuronal lineages in human, mouse and *Drosophila* (Muotri et al. 2005; Coufal et al. 2009; Li et al. 2013; Macia et al. 2017; Chang et al. 2019). Additionally, recent use of an engineered *gypsy* retrotransposon trapping cassette in flies suggested that somatic transposition could also occur in non-neuronal tissues such as the fat body (Jones et al. 2016; Wood et al. 2016) or the intestine (Sousa-Victor et al. 2017). However, a major drawback of using such reporters is that these reporter cassettes could be inactivated by other means than a TE insertion. In addition, the available transgenic lines only report a limited number of TE families. Finally, results obtained with engineered reporters may not necessarily reflect the activity of endogenous elements encoded in the genome.

Genomic sequencing has provided some direct evidence for endogenous somatic retrotransposition though it has almost exclusively focused on the retrotransposition of LINE1 (L1) elements in human cancers (Lee et al. 2012; Solyom et al. 2012; Tubio et al. 2014; Rodić et al. 2015; Doucet-O’Hare et al. 2016; Tang et al. 2017; Rodriguez-Martin et al. 2020) or in human and rodent neuronal tissues (Baillie et al. 2011; Evrony et al. 2012; Upton et al. 2015). However, the first reports of high L1 transposition frequencies in mammalian brains were later shown to be overestimated due to artefacts of sequencing methodology and data analysis (Evrony et al. 2016). Similarly, in *Drosophila*, endogenous somatic TE mobility remains controversial as sequencing performed on populations of adult fly neurons failed to identify true insertions among multiple technical artefacts (Perrat et al. 2013; Treiber and Waddell 2017). Thus, the true extent to which diverse classes of TEs affects genomes of somatic tissues remains to be addressed. Moreover, due to low numbers of somatic insertions recovered thus far from non-cancerous conditions, integration site preferences of TEs in normal tissues *in vivo* are not well understood. Finally, a genetically amenable model system to reliably study somatic transposition is currently lacking.

We have previously established the *Drosophila* midgut as a model system to address the prevalence of somatic mutation in an adult self-renewing tissue (Siudeja et al. 2015). The fly midgut is maintained by a population of intestinal stem cells (ISCs) that divide to self-renew and give rise to two differentiated cell types: absorptive enterocytes (ECs) and secretory enteroendocrine cells (EEs) (Micchelli and Perrimon 2006; Ohlstein and Spradling 2006). Our previous study demonstrated that ISCs acquire genetic mutations including deletions and complex rearrangements, which have important physiological impact on the tissue (Siudeja et al. 2015).

Here, we make use of the fly intestine to demonstrate the contribution of TEs to the somatic genetic variation of an adult tissue. Using whole-genome sequencing of clonally expanded gut neoplasia, we reveal ongoing somatic retrotransposition in the fly midgut. We identify *de novo* TE insertions in the tumor suppressor gene *Notch*, likely contributing to its inactivation and neoplasia formation. Additionally, we apply Oxford Nanopore long-read sequencing of non-clonal healthy adult tissues to provide evidence of tissue-specific differences in retrotransposition. Based on hundreds of high-confidence *de novo* transposition events identified genome-wide, we uncover nonrandom distribution of somatic TE insertion sites in the gut tissue. Transposition occurs throughout the genome and somatic insertions are enriched in genic regions as well as active, enhancer-like chromatin. Overall, by providing direct DNA sequencing-based evidence for *de novo* somatic insertions, we uncover novel features of their *in vivo* biology.

## Results

### Somatic TE insertions in the Notch gene identified in spontaneous intestinal neoplasia

We have previously shown that somatic mutations occur frequently in intestinal stem cells (ISCs) and that the spontaneous inactivation of a tumor suppressor *Notch* in male adult ISCs drives the clonal expansion of mutant cells and formation of gut neoplasia (Siudeja et al. 2015). Neoplasia can be easily distinguished by the clonal accumulation of two intestinal cell types: ISCs expressing Delta and enteroendocrine cells (EEs) marked by Prospero. Our initial sequencing analysis of clonal neoplasia isolated from *ProsGAL4 UAS-2xGFP* (hereafter abbreviated as *Pros>2XGFP*) male flies, revealed inactivation of *Notch* by large deletions or complex genomic rearrangements (Siudeja et al. 2015). In order to expand this analysis and better characterize distinct types of somatic mutations that impact adult ISCs, we generated a large dataset of whole-genome paired-end Illumina sequencing of an additional 30 clonal neoplasia from the same genetic background, as well as four clonal neoplasia from *DeltaGAL4 UAS-nlsGFP* male flies (hereafter abbreviated as *Delta>nlsGFP*), for a total of 37 clonal samples and matched control head DNA sequenced with an average of 47x coverage (***Fig 1A and Supplemental Table 1***). These data are also analyzed by companion paper that addresses structural variation in the same model system (see Methods and also Riddiford, et al, submitted). As expected, a majority of clonal samples showed evidence for inactivation of the Notch pathway by somatic deletions or complex rearrangements (for details see Riddiford et al, submitted). Interestingly, four samples (P15, P47, P51 and D5) did not harbor any other mutation that could explain the clonal expansion, but showed evidence of somatic transposable element sequence inserted in *Notch* (***Fig 1B and C***). Thus, the TE insertions were most likely causative of the clonal expansion and *Notch* mutant phenotype. Strikingly, in sample P15, we observed two integrations within *Notch* (***Fig 1C***), with one of the two events having more sequencing reads supporting the insertion than the other, suggesting that the first insertion inactivated *Notch*, while the second one occurred later during the clonal expansion. All candidate insertions were supported both by clipped-reads mapping partially to a TE and partially to *Notch*, and paired-end reads where one mate-pair is TE anchored and the other is mapped to *Notch* (***Fig 1B***). Among the five candidate insertions identified, three were within the UTR regions of the gene and two TE integrations were in intronic sequences (***Fig 1C***). For all cases described, no read evidence was found for an insertion in the matched head DNA controls. Thus, TE insertions appeared specific for the clonal gut DNA, suggesting they occurred in somatic gut tissue (***Fig 1B***).

**Figure 1.**
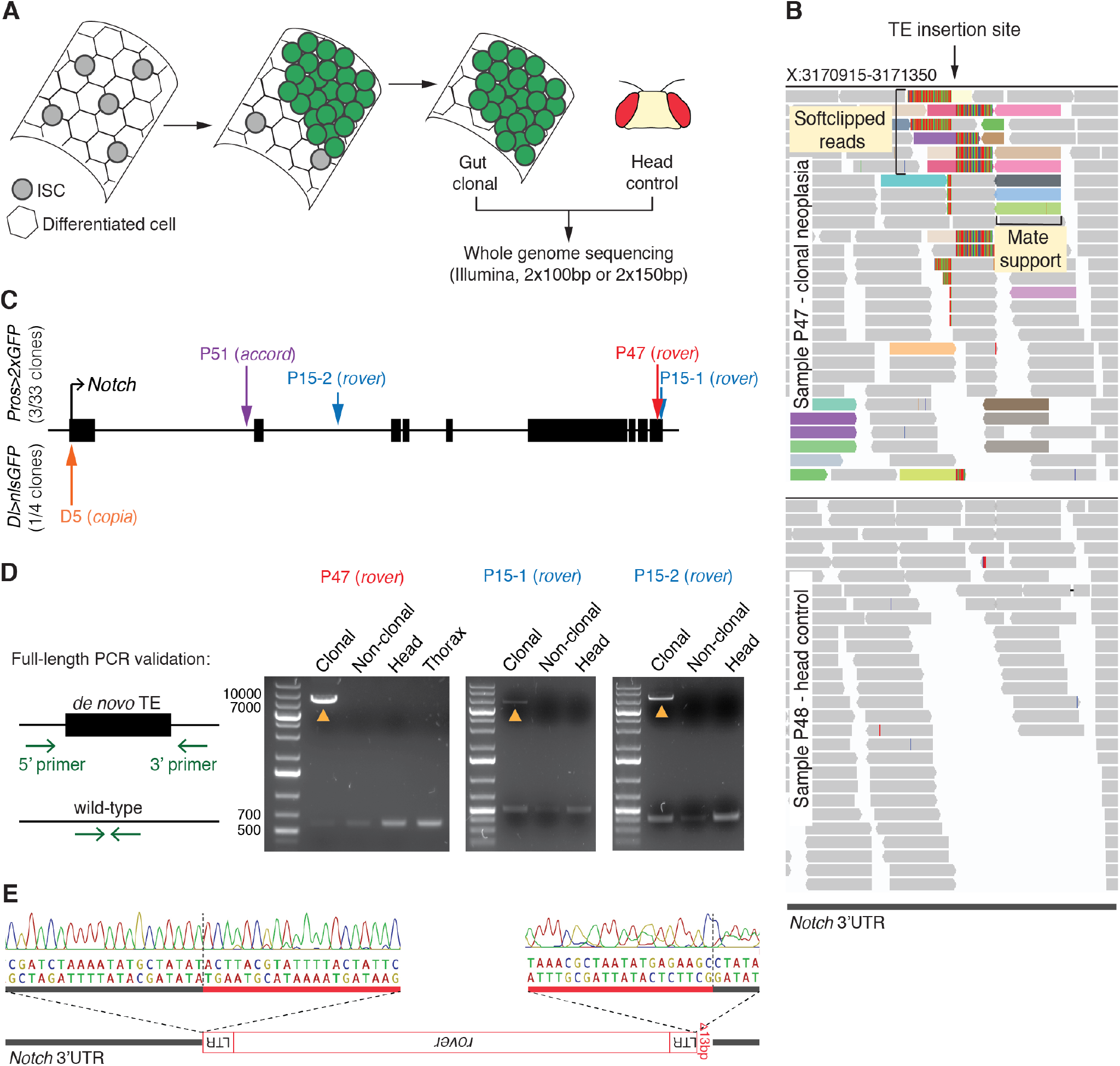
Somatic TE insertions in *Notch* in spontaneous male neoplasia. **(A)** The fly intestine is maintained by the Intestinal Stem Cells (ISCs). Inactivation of *Notch* in a stem cell (in green) leads to a clonal expansion of the mutant cell and neoplasia. The neoplastic gut region was microdissected together with the head of the same fly. DNA isolated from both tissues was subjected to whole-genome paired-end sequencing. **(B)** An Integrative Genomics Viewer (IGV) screenshot of the *Notch de novo* TE insertion site from sample P47 (clonal neoplasia) and its head control, sample P48. Bars represent sequencing reads. Reads supporting the TE insertion are colored according to homology to a specific TE insertion sequence. Multiple colors at a putative insert site frequently indicate homology to different insertions of the same TE family. Two types of supporting reads can be seen: soft-clipped reads - spanning the insertion site and mapping partially to the reference genome and partially to the TE, and mate pair support reads – flanking the insertion site and mapping to the reference genome but with mates (not seen) mapping to the TE. **(C)** The *Notch* locus and the identified somatic TE insertion sites indicated with vertical arrows. Black bars represent exons. Insertions in *Notch* were identified in 3 out of 33 clonal samples from the *Pros>2xGFP* genetic background and in 1 out of 4 *Dl>nslGFP* samples. **(D)** PCR validation of 3 somatic, neoplasia-specific insertions of *rover* elements identified in samples R47 and P15. Primers were designed to target regions flanking the insertion sites. Yellow arrowheads indicate PCR products containing around 8 kb insertion amplified in the clonal DNA but not in the neighboring gut tissue (non-clonal), head or thorax for the same fly. Short wild-type amplicon was detectable in all samples. Thorax DNA sample was not available for sample P15. **(E)** Sanger sequencing of the TE insertion breakpoints in the 3’UTR of *Notch* from sample P47. The *rover* LTR element was inserted in a reverse orientation to *Notch*. The 5’ LTR sequence was truncated by 13 bp. Vertical dashed lines indicate insertion breakpoints. LTR – long terminal repeat

To validate the *Notch* TE insertions, we designed primer pairs flanking the identified insertion sites and performed a full-length PCR amplification using the original genomic DNA as a template (***Fig 1D***). Out of three candidate TE insertions tested, all passed PCR validation, showing ~8 kb insertions likely representing full length TEs. All insertions were amplified only from the clonal neoplastic DNA and not the DNA of matched control tissues from the same fly, confirming that these were true neoplasia-specific somatic TE insertions. Finally, we could further confirm by Sanger sequencing the integration of a *rover* LTR retrotransposon in a reverse orientation in the 3’UTR of *Notch* in the sample P47 (***Fig 1E***).

Altogether, these data revealed that TEs actively transpose in the adult midguts. Importantly, TE insertion can occur into the *Notch* tumor suppressor gene in stem cells, likely driving neoplastic growth in male flies.

### Retrotransposition occurs genome-wide in the fly midgut

Having identified that TEs are mobile in the fly midgut and likely inactivate the *Notch* locus, we then aimed to address the prevalence of somatic transposition on a genome-wide scale. To precisely map somatic TE insertions from our short-read sequencing data, we developed a dedicated pipeline (***Fig 2A***, details in the methods section) and applied it to neoplastic and matched control samples. Briefly, we detected all split-read alignments and discordant alignments supporting TE insertions, isolated clusters of such alignments genome-wide and performed local assembly of the aligned sequence clusters to identify insertion sites with single base-pair resolution. We then filtered for sample-specific calls using all head samples as a panel of normal controls to ensure that somatic, and not germline, insertions are called while retaining the ability to discover recurrent insertions. Finally, we included a manual validation step to exclude any false positives (details in the Methods section).

**Figure 2.**
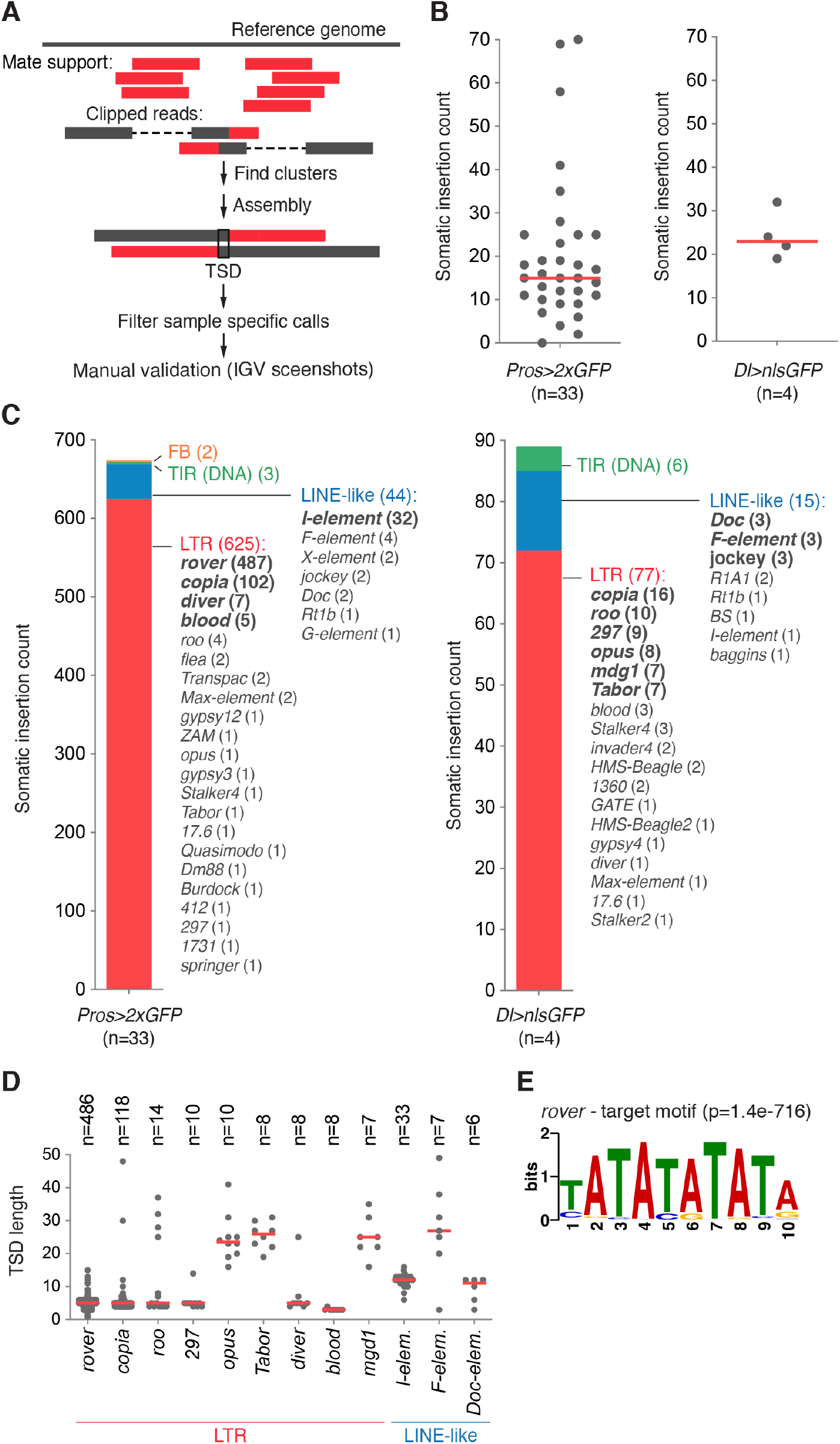
Retrotransposition occurs genome-wide in the fly midgut. **(A)** The bioinformatic pipeline used to identify somatic TE insertions in short-read sequencing datasets. Two types of supporting reads are identified genome-wide: mate support reads, where one of the paired-end reads is mapped to the reference genome, while the other mate (not shown) is associated with a TE, and clipped reads, which span the insertion site and map partially to the reference genome and partially to a TE. Isolated reads were then clustered and assembled to map individual insertion sites. Only insertions with a valid target site duplication (TSD) were retained, sample specific calls were filtered and manual validation of each candidate insertion was performed on IGV. **(B)** The frequency of gut specific somatic insertion sites in the *Pros>2XGFP* and *Delta>nlsGFP* genetic backgrounds. Red lines represent median values. **(C)** The distribution of TE classes active in the two genetic backgrounds studied. TEs were categorized in four main classes: LTR – long terminal repeat retrotransposons, LINE-like – non-LTR retrotransposons, TIR – terminal inverted repeat DNA transposons, and FB – *fodlback* element **(D)** TSD length distribution for somatic insertions of most frequent TE families. Insertions from both genotypes were pooled. Red lines represent median values. **(E)** The target site motif at *rover*-LTR insertions sites recovered from the clonal gut samples.

For further analysis, we retained only insertions bearing a target site duplication (TSD) as a footprint of transposition-dependent events. TSDs are short, identical, duplicated sequences generated on both sides of a TE insertion as a consequence of a staggered endonuclease cut of the target DNA (Feng et al. 1996). We identified a total of 674 (median of 15 per clonal genome) somatic insertions with TSDs from the *Pros>2XGFP* background and 97 (median of 23 per clonal genome) integrations in the *Delta>nlsGFP* samples, all of which were private to gut clonally amplified samples and not present in the matched control DNA, or any of the controls (***Fig 2B, Supplemental Table S1***). Only one gut clonal sample (P13) showed no *de novo* insertions, however this sample was excluded due to low read complexity. In both genetic backgrounds, a great majority of identified insertions were retrotransposons (***Fig 2C***), suggesting that this TE class is the most mobile in the gut tissue. In the *Pros>2XGFP* background the most prominent were insertions of *rover* elements (487 insertions), followed by *copia* (102 insertions), *diver* (7 insertions), *blood* (5 insertions), *roo* (4 insertions), as well as sporadic insertions of other LTR TE families (***Fig 2C***). Among non-LTR retroelements, we identified *de novo* integrations of LINE-like retrotransposons, including 32 *de novo* insertions of *I-elements*, four *F-element* insertions and rare insertions of *X, jockey, Doc, Rt1b, and G-elements*. Insertions of terminal inverted repeat (TIR) DNA elements, such as *1360*, *hobo* or *S-element* (1 integration each) were infrequent (***Fig 2C***). Finally, two somatic integrations of *foldback* elements, belonging to a distinct class of TEs described in *Drosophila* (Truett et al. 1981), were also identified (***Fig 2C***). In the *Delta>nlsGFP* background we mapped 16 insertions of *copia* elements, 10 *roo* integrations, followed by *297* (9 insertions), *opus* (8 insertions), *mdg1* and *Tabor* (7 insertions each) as well as other LTR TE families (***Fig 2C***). Non-LTR LINE-like elements also generated *de novo* integrations of *Doc*, *F, jockey, R1A1, Rt1b, baggins* and *I-elements.* Rare integrations of DNA TIR class TEs, such as *1360*, *pogo, Tc1-2, Tc3* or *S-element*, were also found. Differences in mobile TE families can be noticed between the two genotypes. The most active LTR element family in the *Pros>2XGFP* background, *rover*, as well as LINE-like *I-elements*, were not detected to mobilize in *Delta>nlsGFP* clonal samples. In contrast, *297*, *opus*, *mdg1* and *Tabor* LTR elements that mobilized in the *Delta>nlsGFP* genetic background were rarely detected to mobilize in *Pros>2XGFP* samples. Nevertheless, some TE families, including the second most active element, *copia* LTR, mobilized in both genetic backgrounds. Thus, these data suggest that the repertoire of somatically mobile TEs depends on the genetic background.

To further confirm if the identified TE insertions were indeed true transposition events, we analyzed somatic TSDs for all TE families which produced at least six *de novo* insertions and compared these with known germline TSDs. Most LTR elements generated short TSDs with a median length below 10 base-pairs (5 bp for *rover*, *copia*, *roo*, *297* and *diver*; and 3 bp for *blood*), consistent with TSD lengths reported previously for germline insertions of LTR elements (Dunsmuir et al. 1980; Linheiro and Bergman 2012) (***Fig 2D***). Three LTR elements, *opus*, *Tabor* and *mdg1*, produced unexpectedly long TSDs with a median of 23, 26 and 25 bp respectively, in contrast to 4 bp reported previously for these elements (Linheiro and Bergman 2012). However, with relatively low numbers of somatic insertions of these TE families, it is difficult to conclude if this discrepancy with previously published reports could be biologically relevant. TSDs generated by LINE-like elements were, in general, less strictly defined but centered above 10bp (median of 12, 25 and 11 for *I-*, *F-* and *Doc-elements* respectively, ***Fig 2D***), in agreement with previous reports (Bucheton et al. 1984; Sang et al. 1984; Driver et al. 1989; Berezikov et al. 2000). Finally, we searched for target site motifs for the most represented TEs. Highly significant (AT)-rich target site sequence motif was identified for the *rover* LTR-element reflecting non-random integration (***Fig 2E***). Although there are no previous reports about target site preferences of *rover* elements, TEs from closely-related classes (such as *297 or 17.6)* show similar (AT)-rich target motives (Whalen and Grigliatti 1998; Bowen and McDonald 2001; Linheiro and Bergman 2012). The second most mobile element in our datasets, *copia*, did not show target site preference, which is consistent with previous reports from germline analyses (Dunsmuir et al. 1980). Altogether, our data shows that genome-wide somatic TE integration sites have similar characteristics to germline insertions. This lends further support to the detected TE insertions in the gut being true somatic transposition events, rather than random DNA integrations or products of chimeric reads.

Notably, using our detection criteria, we identified only rare somatic TE insertions in the head samples of both genotypes sequenced (median of 2 insertion/sample in *Pros>2XGFP* heads and 2.5 insertions/sample in *Dl>nlsGFP* heads, ***Supplemental Fig S1A and B***). However, the frequency of transposition between gut and head samples cannot be directly compared in this assay. Indeed, the head is a heterogeneous cell population, therefore somatic transposition in a few cells of the head would be below the detection level in our analyses. In contrast, the intestinal neoplasia are clonal expansions of single ISC genomes, increasing likelihood of detecting TE insertions. Accordingly, the rare somatic insertions identified in head samples had only a few clipped and mate-pair supporting reads, reflecting that these were likely rare events present in limited numbers of cells (***Supplemental Fig S1C***). This difficulty to detect TE insertions in non-clonal fly head DNA is also in agreement with recently published data (Treiber and Waddell 2017). Because single cell-insertions are unlikely to be detectable in our assay, we believe that the identified head insertions occurred likely during brain development leading to a small clone of cells harboring the TE insert, rather than in an adult fly brain, which is post-mitotic. Alternatively, they could represent rare but recurrent insertions arising independently in multiple cells of the adult fly brain.

Overall, we conclude that somatic retrotransposition in the fly midgut is not limited to the *Notch* locus, but occurs genome-wide. LTR-elements are the most active, while LINE-like retrotransposons mobilize less frequently. Although TE families identified as the most mobile can differ between fly strains, our data suggests that retrotransposons are frequently active in gut tissue.

### TE insertions arise before and after the clonal expansion

To better understand when somatic transposition occurs in the fly gut, we then used allele frequency estimates to time genome-wide *de novo* integrations identified in clonal samples relative to the event inactivating *Notch* and initiating the clonal expansion (***Fig 3A***). The allele frequency is the ratio of sequencing reads supporting and opposing any given insertion. A TE insertion could arise before the onset of neoplasia, either during development or in the young adult gut, and be present in some cells of the normal tissue. Upon the *Notch* pathway inactivation, a stem cell would initiate clonal expansion and, at the time of analysis, the insertion would be present in all clonal cells as well as neighboring “normal” cells isolated for sequencing along with the clone. Thus, assuming the observed allele frequency represents the true population allele frequency, such a variant would present estimated allele frequency equal or greater than the *Notch* mutation. Alternatively, transposition could also occur after the clonal expansion, in which case such insertion would then be present in a fraction of cells of the clone and show estimated allele frequency lower than that of the Notch pathway inactivating mutation.

**Figure 3.**
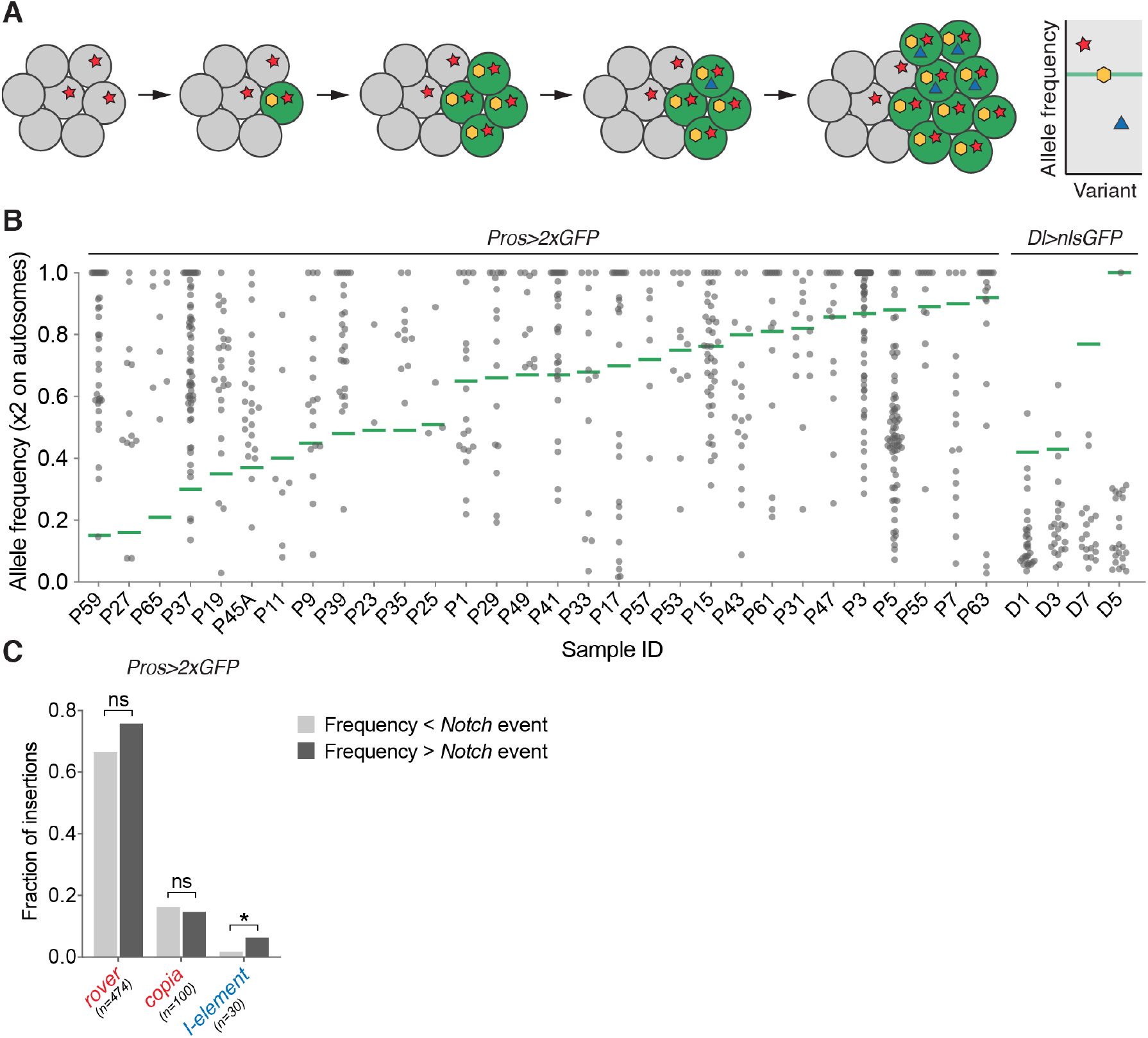
Somatic retrotransposition occurs before and after the clonal expansion. **(A)** A somatic insertion may arise in a normal tissue and be present in a fraction of cells of the tissue (red star). If a second somatic event (yellow) inactivates the Notch pathway, the mutant cell (indicated in green) will initiate the clonal expansion amplifying somatic variants already present in its genome. Finally, any insertion that occurs after the clonal expansion (blue triangle) will be present in a subset of neoplastic cells. The graph represents theoretical allele frequencies of somatic variants obtained from the sequencing of clonal tissue samples. The allele frequency of a *Notch* inactivating event, marking the onset of neoplasia is represented with a green horizontal line. A somatic insertion with allele frequency higher than the *Notch* inactivating event was likely present in the tissue before the clonal expansion. In contrast, an insertion with allele frequency lower than the *Notch* inactivating event, likely occurred after the initiation of neoplasia and is thus subclonal. **(B)** The calculated allele frequencies for all somatic TE insertions identified in neoplastic samples (*Pros>2GFP* and *Dl>nlGFP* genetic backgrounds). The onset of neoplasia is represented with a green horizontal line. Three samples (P45B, P51 and P21), where timing of the neoplasia onset could not be unambiguously estimated, were excluded from this analysis. **(C)** The distribution of somatic insertions with frequencies higher or lower than *Notch*-inactivating mutations for the most represented LTR-elements (*rover* and *copia*) and LINE-like *I*-elements. ns- not significant, *- p<0.01 (Fisher’s exact test, two-tailed), n=number of insertions

In both genetic backgrounds sequenced, we uncovered TE insertions causative for *Notch* inactivation and initiating neoplasia formation, indicating pre-clonal TE mobility (***Fig 1C***). In the *Pros>2xGFP* background 63.5 % of genome-wide somatic integrations showed allele frequency higher than the *Notch* pathway inactivating event in the same sample (***Fig 3B***). These insertions were thus likely present in the tissue before the onset of neoplasia and occurred either during gut lineage specification in development or in young adult life. The remaining insertions (36.5 %) were of lower allele frequency than *Notch* pathway inactivating events, indicating that transposition continued to occur within clonal populations of cells after the initiation of a neoplasia (***Fig 3B***). High- and low-allele frequency insertions were detected for all TE families for which insertion counts were high enough to allow such analysis (***Fig 3C***). The LTR-elements *rover* and *copia* inserted equally often before and after the neoplasia initiating event. In contrast, LINE-like *I-element* integrations were moderately, but significantly, enriched among variants with allele frequencies higher than the *Notch* events, suggesting that this TE family might be more active during development in the precursor lineage of the gut or in the normal adult guts prior to the onset of neoplasia. In contrast to *Pros>2XGFP* neoplastic clones, insertions in *Delta>nlsGFP* samples were largely subclonal to the neoplasia initiating events (96.8 %), indicating that in this genetic background a majority of detected TE integrations occurred in the adult gut after the onset of neoplasia (***Fig 3B***). However, the remaining few insertions, including one in *Notch*, likely causative of *Notch* inactivation, indicate that pre-clonal mobility also occurred in this genetic background.

Together, these results imply that somatic retrotransposition in the fly gut can drive inactivation of a tumor suppressor *Notch*, and that it can occur before and after the clonal expansion. This suggests that retrotransposon activity is not restricted to neoplasia and can act through adult life in the gut and perhaps also during development.

### TE expression levels do not predict their mobility

The expression and activity of LINE1 (L1) elements, the only somatically active TEs identified to date in the human genome, are increased in different tumor types (reviewed in (Burns 2017)). Our allele frequency analysis suggested that many TE insertions preceded clonal expansion, suggesting that TEs were active in a normal tissue. Nevertheless, we wished to address whether the initiation and clonal expansion of gut neoplasia could lead to transposable element deregulation. Thus, we asked if inactivation of *Notch* in a stem cell, which leads to a clonal neoplasia, could also cause increased TE expression. To do this, we compared TE expression levels in previously published RNA sequencing data of FACS-sorted wild-type and *Notch* RNAi knockdown ISCs (Patel et al. 2015). Expression of most TEs was not affected upon *Notch* knock-down (***Supplemental Fig S2***). Five TE families were significantly upregulated and four families were downregulated in *Notch* knock-down cells. However, changes were of low magnitude (fold change ranging from 1.5 to 3.5) and none of the differentially expressed TEs in *Notch* RNAi ISCs overlapped with the mobile TE classes identified in our assays (***Supplemental Fig S2***). Even though this data set comes from a different genetic background than the one we used to isolate neoplastic clones, the results suggest that neither inactivation of the Notch pathway nor hyperproliferation of the gut tissue are sufficient to strongly deregulate TE expression.

We next asked whether there was a correlation between expression levels of TE classes and their mobility. To do so, we performed RNA sequencing of normal (non-neoplastic) *Pros>2xGFP* midguts and compared this with our data on *de novo* TE insertions (***Fig 4A***). Most TEs showed very low (TPM, transcript per million< 1) or low (1 < TPM < 5) transcript levels. Eight TE families (7 LTR and 1 LINE-like element) showed TPM values above 5. The most highly expressed elements, *copia-*LTRs, contributed 73.5 % of all TE-mapping transcripts (TPM=481), while the second and third most highly expressed TEs, *springer* and *accord*, represented 6.3 % and 2 % respectively. *copia* was also the second most mobile element identified in this genetic background. Strikingly, mobility did not correlate with TE expression levels, as the most active LTR-elements *rover* as well as the most active LINE-like *I-elements*, were both very lowly expressed in the tissue (TPM<1) (***Fig 4A***). For certain moderately expressed TE families, such as LTR element, *invader1*, or the LINE-like element, *Juan*, the canonical TE sequence was only partially covered, suggesting that full length, transposition-competent copies were not transcribed (***Fig 4B***). Notably, these data show that, at the tissue-wide scale, steady-state levels of TE transcripts are not good predictors of TE mobility.

**Figure 4.**
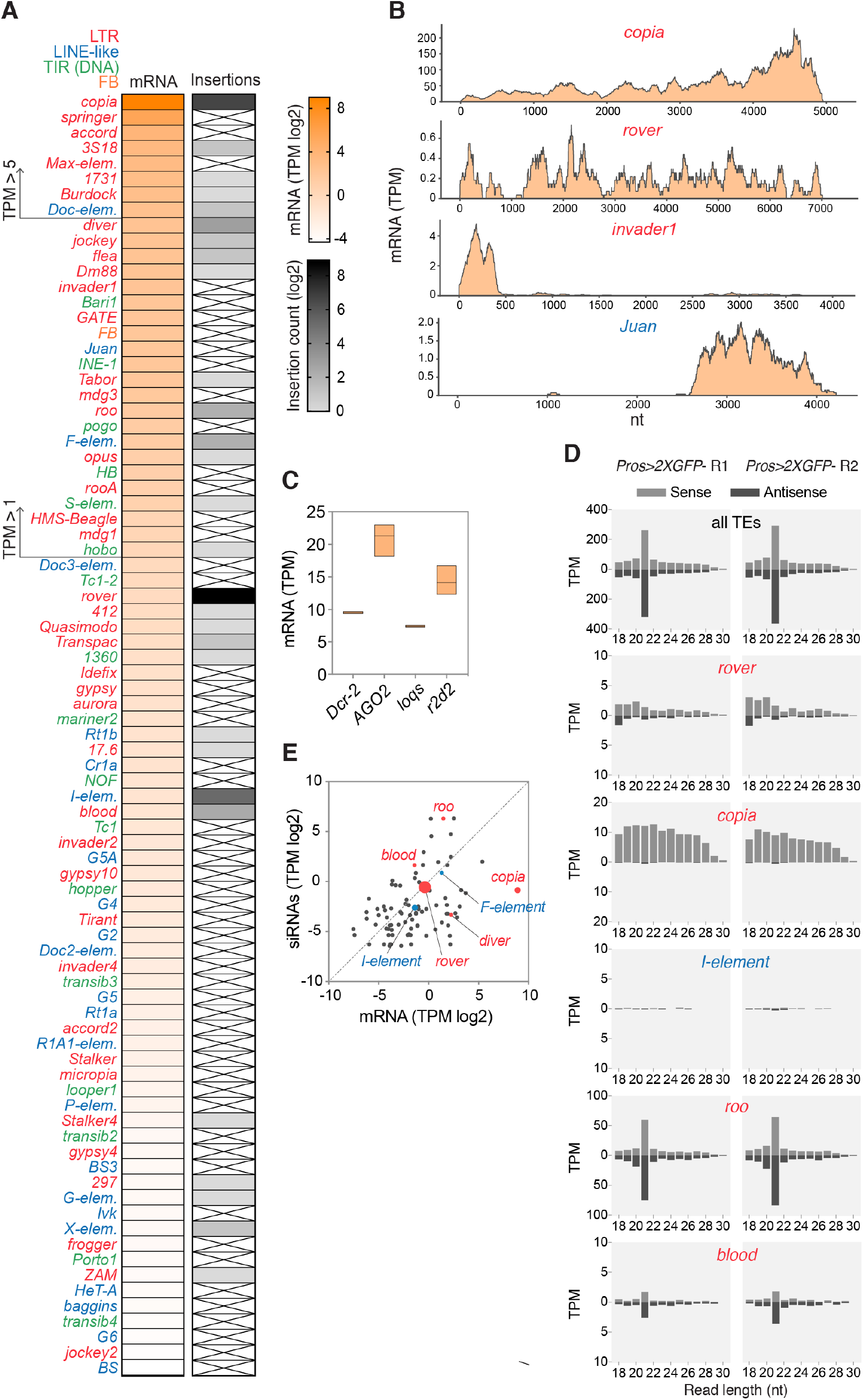
TE expression and siRNA pathway activity in the fly midgut. **(A)** Heatmaps representing normalized TE expression levels (in log2(TPM), transcripts per million) and mobility (log2(insertion counts)) in *Pros>2xGFP* midguts. TEs with TPM values below 0.05 (log2(TPM)< −4.3) are not depicted. Crossed out cells represent no somatic insertions of that family identified. **(B)** Normalized read coverage over the full-length canonical sequence of selected TE families. **(C)** Normalized expression levels of siRNA pathway genes. Bars represent the minimum, the maximum and the mean from three biological replicates. **(D)** The size distribution of sense and antisense reads from gut small RNA fractions mapping to all TEs (upper panel) or selected TE families mobilizing in the gut. R1 and R2 are two biological replicates. **(E)** Scatter plot of normalized transcript (mRNA) levels and antisense, 21nt siRNA levels for all TE families. Transposons generating somatic insertions are highlighted in red (LTR-elements) or blue (LINE-like), with a symbol size reflecting the somatic insertion counts.

On the post-transcriptional level, somatic control of TEs is mostly achieved by the endogenous siRNA (endo-siRNA) pathway (Chung et al. 2008; Czech et al. 2008; Ghildiyal et al. 2008). Our gut transcriptome analysis showed that siRNA pathway genes, *Argonaute 2* (*AGO2*), *Dicer-2* (*Dcr-2*), *loquacious* (*loqs*) and *r2d2* were expressed, suggesting that this pathway is functional in the fly midgut (***Fig 4C***). This was further confirmed by sequencing of the gut small RNA fraction, which detected short 21-nucleotide sense and antisense reads complementary to TEs, as expected for the *Drosophila* siRNAs (***Fig 4D***). We found that siRNA levels were not directly proportional to the TE transcript levels (***Fig 4E***). The five active TE families (*rover*, *copia*, *I*-element, *diver* and *F*-element), responsible for 93.8% of all insertions, showed low levels of 21-nt antisense siRNAs. Thus, post-transcriptional silencing by siRNAs of these elements could be inefficient. Nevertheless, low siRNA levels were not a prerequisite for mobility, as we also detected *de novo* insertions of TEs (including *blood* and *roo*) that had abundant siRNAs present (***Fig 4D and E***). Importantly, this implies that low levels of siRNAs could allow for the somatic mobility of some TEs, while other TEs retain their ability to mobilize even in the presence of abundant siRNAs.

Recent reports suggested that the piwi-interacting RNA (piRNA) pathway, known to control TEs in the gonads (Brennecke et al. 2008; Chambeyron et al. 2008), could also play a role in somatic TE silencing (Perrat et al. 2013; Jones et al. 2016; Sousa-Victor et al. 2017). However, it remains to be proven whether piRNAs are indeed produced in somatic tissues. We did not detect abundant 23-30 nucleotide-long RNAs (characteristic of piRNAs) in the analyzed gut small RNA samples (***Fig 4D***). Thus, if piRNAs are produced in the gut, they are at low levels and were under our detection limit. In contrast, 23-30 nucleotide-long RNAs with a typical “ping-pong” signature, indicative of piRNAs (Brennecke et al. 2007; Gunawardane et al. 2007), were easily detected in ovary controls of the *Pros>2xGFP* females. These were complementary to all TEs, including somatically active TE families (*rover*, *copia*, and *I-element*), suggesting that piRNA-mediated TE silencing of these TEs was properly established in the female germline (***Supplemental Fig S3A and B***). There were also no significant differences in ovary piRNA levels between two parental stocks (*ProsGAL4* and *UAS-*2xGFP) used to obtain the *Pros>2xGFP* flies (***Supplemental Fig S3C and D***). Thus, the observed somatic TE activity could not be explained by differences in active TEs between the parental genotypes, as previously documented in *Drosophila* dysgenic crosses (Brennecke et al. 2008).

Altogether, we find that in the fly gut neoplastic transformation is not necessary for TE expression and that many TE families are transcribed in the normal gut tissue. At the tissue-wide scale, TE RNA levels do not correlate with somatic mobility and even very low transcript levels can be sufficient for active transposition. Although post-transcriptional control by the siRNA pathway is in place, some retrotransposons escape this control and mobilize in the tissue.

### Tissue-specific transposition

To further address TE mobility in normal tissues without a clonal expansion, we decided to apply long-read sequencing to bulk genomic DNA obtained from either pooled midguts or pooled heads from the same individuals of the *Pros>2xGFP* background. High molecular weight genomic DNA was sequenced to 85x coverage using the Oxford Nanopore Technology (ONT). We then detected all full-length, non-reference and tissue-specific TE copies entirely contained in a sequencing read (***Fig 5A***). Considering that in the absence of clonal expansion, any somatic insertion would be very rare in sequencing of bulk DNA, we extracted only insertions supported by a single read (“singletons”) and classified them as potentially somatic. To help to exclude germline variants, we eliminated all insertions detected in both gut and head DNA pools. Finally, as in the short-read sequencing datasets, we retained only insertions generating a valid TSD, a footprint of active transposition. Among all singleton insertions, *rover* LTR-elements was detected the most frequently in gut DNA (152 insertions) (***Fig 5C***). Importantly, the fact that *rover* singleton insertions are the most frequent in both the long-read data as in the Illumina short-read data, supports the notion that these are likely somatic *de novo* insertions. Additionally, mapped *rover* insertions found in long-read ONT data had identical AT-rich target site motifs to those identified in clonal samples with Illumina sequencing (***Fig 2E and 5C***). Similar to the short-read Illumina sequencing, singleton reads from the gut were also found containing other LTR-elements (such as *springe*r, *copia, roo*), LINE-like elements (*Doc-*, *I-element*) and DNA TIR elements (*hobo*, *Bari1, S-element*) (***Fig 5C***). While a total of 191 singleton insertions, representing putative *de novo* integration events, were recovered from the gut DNA, 24 singletons were also found in the head DNA. This suggests that somatic TE mobility occurs both in the gut as well as in the head, further supporting our findings of rare inserts in the head from the short-read sequencing data (see ***Supplemental Fig S1***). Interestingly, with the exception of singleton insertions from *roo* LTR-elements, almost all singletons from other TE families were specifically found in one of the two tissues, suggesting that TE family activity may be tissue-specific. Singletons from LTR-elements *rover*, *springer* and *copia*, contributing together 82.8 % of all putative somatic insertions, were found only in the gut tissue. Singletons of other elements (such as *Stalker2*), although rarer, appeared specific to the head DNA (***Fig 5C***). This suggest that the repertoire of somatically active TEs may differ between different tissues.

**Figure 5.**
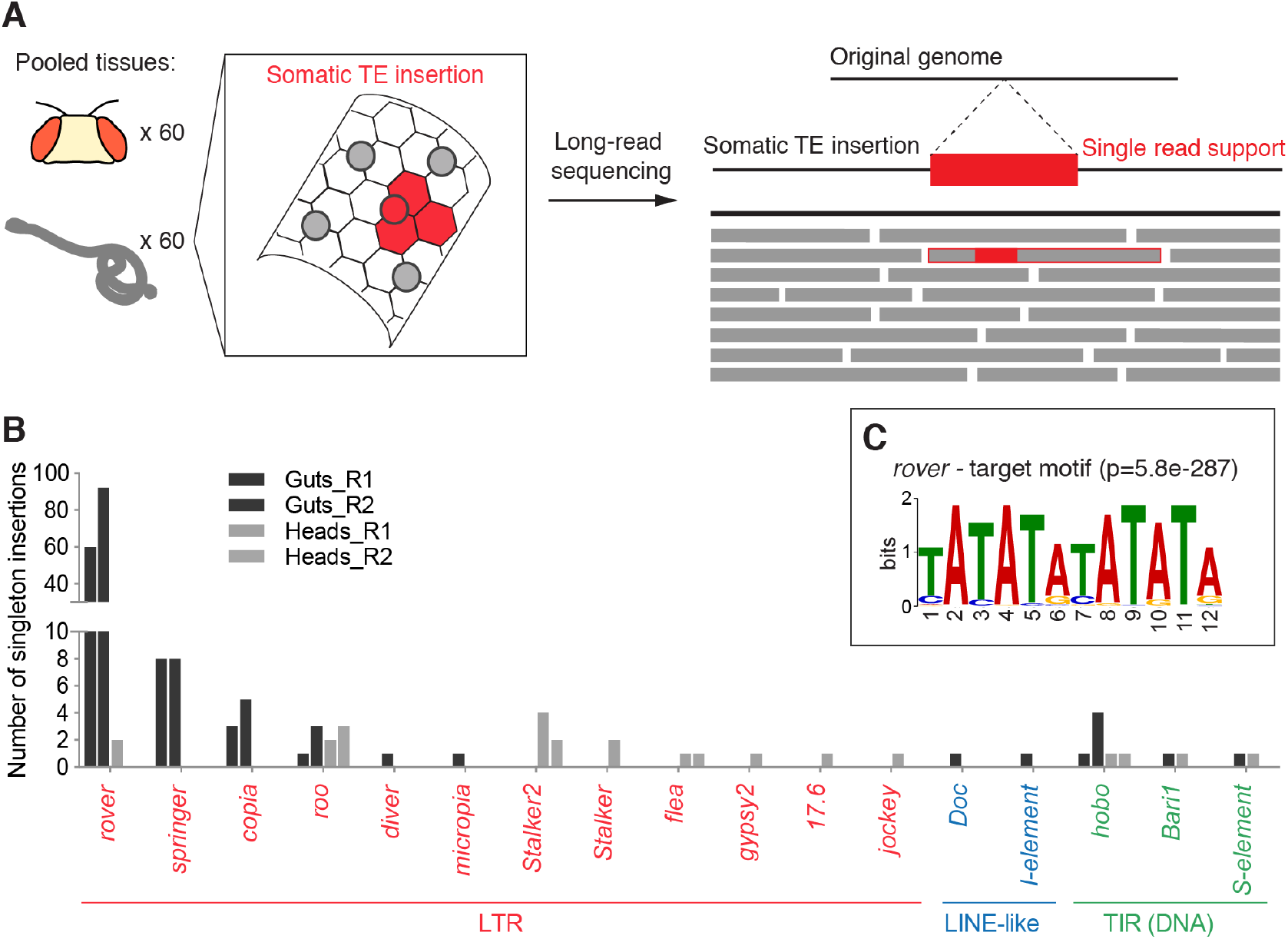
Long-read sequencing implies tissue specific transposon mobility. **(A)** Genomic DNA was isolated from pools of 60 guts or 60 heads dissected from the same individuals and subjected to ONT long-read sequencing. If a somatic insertion occurs in a tissue and does not undergo clonal expansion, it will be present in a small fraction of cells within the pool sequenced. Long-read sequencing allows to identify putative somatic integrations as rare TE inserts, fully contained in a single continuous sequencing read and generating a valid TSD. **(B)** TE families with tissue-specific, singleton TSD-bearing insertions detected in pooled gut or head DNA. R1 and R2 are two biological replicates. **(C)** The target site motif at gut-specific *rover* singleton insertion sites identified with the long-read sequencing.

To gain further insight into the tissue-specificity of transposition, we then asked if the same TEs were mobile in the germline, leading to heritable *de novo* insertions. To do so, we sequenced 18 individual flies from the progeny of *Pros>2xGFP* parents (***Supplement Figure S4A***) and detected *de novo* germline insertions present in any of the progeny and not in the parents. We discovered 3 *de novo foldback* element insertions. Of note, we found no germline *de novo* insertions of any of the TE families found active in the somatic cells of the gut. This was in agreement with the analysis of small RNA fractions from ovaries indicating that somatically mobile TEs were properly silenced in the germline (***Supplement Fig S3A and B***).

Altogether, with the use of long-read sequencing, we provide further evidence supporting active transposition in the normal fly intestine. Furthermore, our data comparing gut and head DNA as well as inherited *de novo* insertions, indicates that 1) distinct TEs can be active in the different somatic tissues and 2) active TEs may differ between the soma and the germline.

### Somatic transposition is enriched in genes and regions of active enhancer-like chromatin

The identification of hundreds of somatic TE insertions with base-pair resolution, allowed us to address whether somatic transposition acted uniformly throughout the genome or specifically affected particular genomic regions. To shed light on preferential landing sites of somatically active TEs, we analyzed genomic and chromatin features of the *de novo* somatic TE insertion sites.

Analyzing the genome-wide distribution of somatic TE insertions from the clonal gut samples in mappable regions of the genome revealed that integrations occurred broadly across *Drosophila* chromosomes (***Fig 6A***). Importantly, somatic transposition was very frequent in genic regions. 78.5% of all candidate insertions fell inside annotated genes and an additional 5.6% targeted potential promoter sequences less than 1kb upstream of genes. Although they were depleted from coding sequences, somatic TE integration sites were significantly enriched in introns and 3’UTRs (***Fig 6B***). Similar genic preference was evident in the singleton insertions identified by the long-read sequencing of normal gut DNA pools (***Supplemental Fig S5***), suggesting that it was not influenced by the clonal expansion. 82.3% of all singleton insertions fell inside annotated genes and additional 6.2% in gene promoters. Importantly, in both data sets, we found insertions within or nearby (<500bp upstream of the TSS) genes with established roles in the regulation of gut and ISC homeostasis (***Fig 6C, Supplemental Table S2 and S3***). Apart from the insertions in *Notch*, which were selected for in clonal samples, we detected insertions in genes involved in the EGFR (*pointed, EGFR*), JNK (*puckered*), JAK/STAT (*Stat92E*), Wnt (*frizzled*), insulin (*Insulin-like receptor*) and VEGF (*Pvf1-3)* pathways, as well as chromatin modifiers (*kismet*, *osa)* and other regulators of ISC homeostasis. Some of the affected genes were hit multiple times (***Fig 6C, Supplemental Table S2 and S3***). It is possible that these genes are hot-spots for TE insertions, perhaps due to genome sequence or structure. Alternatively, these insertions could drive positive selection of the resulting cell lineage through promoting stem cell proliferation, resulting in their post-insertion enrichment. Thus, we find that somatic TE insertions are enriched in genic regions and frequently target genes, including those with important functions in the gut.

**Figure 6.**
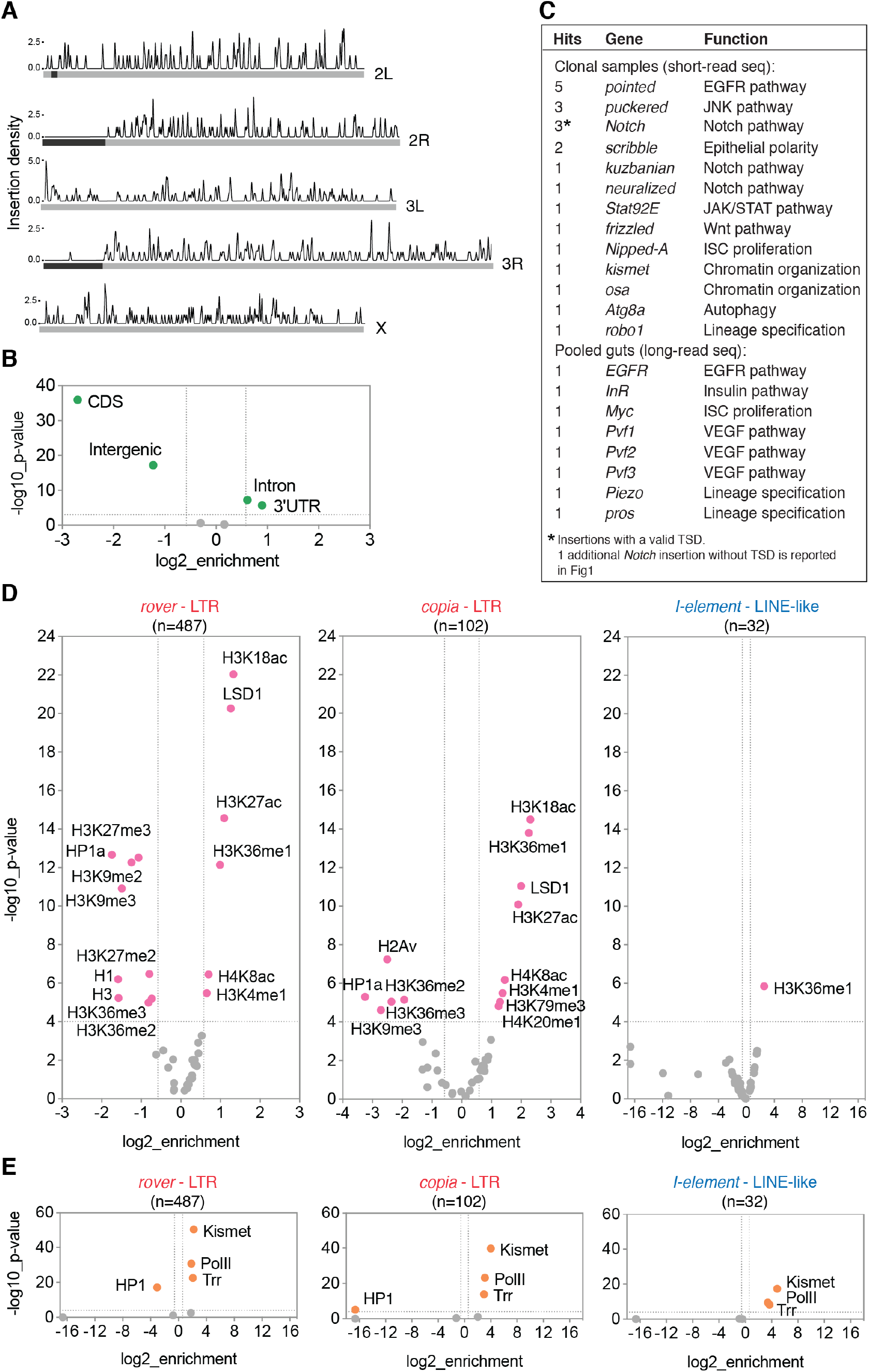
TEs frequently insert in genes and regions of open enhancer-like chromatin. **(A)** The distribution of somatic TE insertion sites on the *Drosophila* chromosomes. Dark grey boxes represent heterochromatic regions. **(B)** Somatic TE insertion sites were depleted from intergenic and exonic sequences and enriched in introns and 3’UTR regions of the fly genome. **(C)** Selected genes relevant for the gut physiology with putative somatic insertion sites. Genic regions +/− 500bp were considered. **(D)** Correlations of somatic insertion sites of the three most represented TE families (*rover*, *copia* and *I-element*) with modENCODE tracks for adult fly tissues. **(E)** Correlations of somatic insertion sites of the three most represented TE families (*rover*, *copia* and *I-element*) with DamID tracks for adult fly intestinal stem cells (ISC). Colored data points and labels highlight significant positive or negative correlations (p<0.0001, −1.5>enrichment>1.5). *de novo* insertions from the *Pros>2XGFP* clonal gut samples obtained with the short-read sequencing were used for all plots.

To further probe into integration preferences of somatic TE insertions, we investigated the overlap between candidate integration sites mapped in clonal (***Fig 6D***) and pooled gut samples (***Supplement Fig S5B***) with publicly available *Drosophila* modENCODE datasets profiling chromatin features and transcription factor binding sites (The modENCODE Consortium et al. 2010). Comparing clonal gut TE insertion sites of LTR-elements (*rover* and *copia*) with tracks from adult fly tissues revealed significant depletions from genomic regions enriched in silent chromatin features, such as methylated histone H3 (H3K9me2/3, H3K27me3), Heterochromatin Protein 1a (HP1a) or linker histone H1 (Riddle et al. 2011). Although less strongly, LTR-element insertion sites were also depleted from regions marked by H3K36me2/3, typically associated with exons of transcribed genes (Kharchenko et al. 2011). This negative correlation is agreement with the depletion of TE insertion sites from coding regions as documented above (***Fig 6B***). In contrast, we observed a strong positive correlation between LTR-element integration sites and genomic regions rich in acetylated histone H3 (H3K18ac, H3K27ac) and H4 (H4K8ac) as well as H3K36me1, marks associated with active promoters, transcribed regions and enhancers (Kharchenko et al. 2011; Nègre et al. 2011). *De novo* insertions were also significantly enriched in genomic regions bound by LSD1/Su(var)3-3, a histone lysine-demethylase responsible for removing histone H3K4-methyl marks from active promoters (Shi et al. 2004; Stefano et al. 2007). The correlations for LTR-element insertions identified in clonal and pooled DNA samples were very similar (***Fig 6DB* and *Supplement Fig S5B***), suggesting that the distribution of somatic insertion sites was the same for normal and neoplastic tissue. Correlations of the LINE-like *I-element* insertion sites were much weaker than those obtained for LTR elements, and should be interpreted with caution due to a low total number of insertions (***Fig 6D***). However, the enrichment in H3K36me1 was also significant for this non-LTR TE family. Consistent with the insertion timing analysis implying that transposition could act pre-clonally and during development (***Fig 3B***), comparable enrichments and depletions were also found for short-read clonal (***Supplement Figure S6***) and long-read bulk (***Supplement Figure S5B***) gut sequencing data sets with the modENCODE tracks derived from *Drosophila* larval stages.

To confirm whether similar TE insertion site enrichment could be observed with gut-specific chromatin features, we used our recently published DamID profiles of chromatin factors in intestinal stem cells (Gervais et al. 2019) (***Fig 6E***). In agreement with the results obtained with the modENCODE datasets, somatic TE insertions from clonal gut samples were highly enriched in genomic regions bound by chromatin modifiers Kismet and H3K4 monomethyl-transferase Trithorax-related (Trr) as well as RNA polymerase II (Pol II) (*rover*, *copia* and *I-element*). All these factors were previously shown to map open, transcriptionally active chromatin (Marshall and Brand 2017; Gervais et al. 2019). Concurrently, somatic transposition sites of LTR-elements were strongly depleted from repressed chromatin domains bound by HP1 (a reader of histone H3K9me3) (***Fig 6E***). Finally, enrichment in chromatin domains bound by Kismet and Pol II, and depletion in HP1-bound sites were also significant for putative somatic singleton insertions identified in non-clonal gut DNA pools (***Supplement Fig S5C***), suggesting that the insertion site enrichment in active chromatin also occurred in normal tissues without clonal expansion.

Taken together, our data revealed non-random distribution of retrotransposon insertions sites in a somatic tissue *in vivo*. Somatic transposition more frequently affects genic regions of the genome and is enriched in open, transcriptionally active chromatin.

## Discussion

Our study provides a genome-wide view of how transposable elements mobilize in a somatic tissue. We show that endogenous retrotransposons mobilize in the fly gut and create *de novo* insertions genome-wide. Somatic insertions preferentially affect genes and open chromatin, and they can occur in the *Notch* tumor suppressor, likely leading to clonal neoplasia.

### Somatic retrotransposition in the fly intestine

By whole-genome sequencing of clonally expanded gut neoplasia, we were able to detect hundreds of high confidence retrotransposition events. Clonal expansion brings an advantage of amplifying *in vivo* any genetic variant otherwise present in a tissue with a very low frequency. A similar approach has previously been taken to demonstrate somatic transposition in human tumor samples. Indeed, retrotransposition was observed in many tumor types, including lung, ovarian, breast, colorectal, prostate, liver, pancreatic, gastric, esophageal and hepatic cancers (Iskow et al. 2010; Lee et al. 2012; Solyom et al. 2012; Shukla et al. 2013; Helman et al. 2014; Tubio et al. 2014; Doucet-O’Hare et al. 2015; Ewing et al. 2015; Paterson et al. 2015; Rodić et al. 2015; Scott et al. 2016; Schauer et al. 2018). Most of these studies failed to detect somatic integrations in matched healthy tissues, leading to the conclusion that transposition was likely limited to the cancerous state. However, careful analysis of gastro-intestinal and esophagus tissues did suggest that active retrotransposition occurring in normal cells, can undergo clonal expansion in a tumor context (Ewing et al. 2015; Doucet-O’Hare et al. 2016). Likewise, here we provide evidence that TEs are expressed and mobilize in a normal fly gut, without a prerequisite of neoplastic transformation. Indeed, our analysis of variant allele frequencies in clonal datasets suggests that transposition can occur in the adult tissue both in normal cells prior to the clonal expansion and in neoplastic cells undergoing uncontrolled proliferation. Studies of the mammalian brain have provided different pieces of evidence suggesting that somatic transposition can occur in the embryo, during neurogenesis as well as in mature neuronal cells (Evrony et al. 2012, 2015; Upton et al. 2015; Faulkner and Garcia-Perez 2017). Likewise, we detect TE insertions with variant frequencies higher than the event driving clonal expansion but with no supporting evidence in the head DNA, suggesting that somatic TE insertions could also be acquired during development and gut lineage specification as well as during adult life.

### Applying long-read sequencing to detect somatic transposition

We provide further strong evidence for somatic transposition acting in a normal tissue with the use of long-read sequencing technology to assess rare, single read support insertions representing likely *de novo* TE integrations in bulk tissue DNA in the absence of clonal expansion. Indeed, this technology offers important benefits over classical short-read sequencing in mapping non-referenced TE insertions. It enables full-length detection of inserts within one sequencing read, resolving not only both ends but also the entire length of an insert. Moreover, it outperforms short-read technology in the analysis of low-complexity genomic regions where short-read mapping poses particular difficulty. Thus, rare somatic TE insertions can be detected from pooled DNA libraries, provided that robust controls are implemented to help to exclude germline variants in a population. Alternatively, the use of single individuals, rather than pooled tissues, would rule out germline variants. Thus, by the use of long-read sequencing we put forward a novel methodology for detecting somatic TE activity. A similar approach has very recently been proposed to perform epigenomic profiling and non-referenced TE mapping in human datasets (Ewing et al, bioRXiv 10.1101/2020.05.24.113068), further showing that long-read sequencing will certainly gain on popularity in the field. As our results obtained with this technology are highly consistent with our results obtained with short-read sequencing of neoplastic clones, we believe the singleton reads detected with long-read sequencing are very likely *de novo* somatic events. Thus, we reveal ongoing somatic transposition in the gut tissue using two complementary genomic approaches.

### Transposition across different tissues

Historically, focus has been on somatic transposition in neuronal tissues, where TE mobility was proposed to contribute to functional differences between individuals (Erwin et al. 2014). Here, with the use of long-read sequencing, we detected putative somatic TE insertions in the gut and the head tissues. Our data suggest that TE families active in the gut are largely not mobile in the head of the same individuals, implying that the repertoire of mobile TEs might vary between different somatic tissues. Indeed, many studies have begun to uncover tissue- and cell-specific patterns of TE expression in human and model organisms (Mietz et al. 1992; Faulkner et al. 2009; Philippe et al. 2016; Deininger et al. 2017; Pehrsson et al. 2019; Chung et al. 2019; Ansaloni et al. 2019; Sanchez-Luque et al. 2019) (Treiber and Waddel, bioRxiv 10.1101/838045). However, in most cases, the lack of data on somatic insertions hinders the direct comparison between transcriptional activity and mobility. Importantly, here we show that in the gut tissue, TE transcript levels do not parallel insertional activity. However, we cannot exclude that active TEs could be expressed in a cell-type specific manner and that cell-type specific TE expression patterns could correlate better with the mobility. Nevertheless, our data suggest that caution should be taken when using transcript levels as a proxy for TE insertional activity. How additional factors, aside from those regulating RNA transcript levels, may contribute to tissue-specific somatic TE mobilization, remains to be determined.

Apart from gut vs head specificity, we also show that somatically active TEs were not detected to be mobile in germ cells. Thus, our data speaks against an overall deregulation and high retrotransposon activity, as previously documented in the germline of *Drosophila* dysgenic crosses (Kidwell et al. 1977; Pélisson 1981; Rubin et al. 1982) or in other rare genetic backgrounds with bursts of TE activity in the germline and soma (Gerasimova et al. 1985; Georgiev et al. 1990).

### TE insertions distribute non-randomly in the somatic genomes

Mapping somatic retrotransposition insertions with base-pair resolution reveals enrichments in insertion site distribution of endogenous retrotransposons *in vivo*. Indeed, we show that the distribution of TE insertions is not random, but influenced by the genome sequence composition as well as by the local chromatin state. Gut retrotransposon insertions are enriched in transcriptionally active, enhancer-like chromatin, a bias that is similar to the insertion site preferences previously observed for the murine leukemia virus (MLV) and the PiggyBac transposon in human T-cell cultures (Gogol-Döring et al. 2016; Sultana et al. 2017, 2019). Similarly, recent analysis of TE insertion site preferences in human cancer genomes revealed enrichment in DNase hypersensitive open chromatin and depletion in histone H3K9me3-rich heterochromatin (Rodriguez-Martin et al. 2020). Our data uncover similar insertion site enrichments *in vivo*, not only in the context of neoplastic clones but also in a normal tissue. In the fly gut, as transposition acts in the context of renewing and dividing tissue, the uncovered insertion site distribution is likely a result of pre-insertion target site choice as well as post-insertion selection in the tissue, as previously demonstrated for *de novo* L1 insertions in human culture cells (Sultana et al. 2019). Negative selection probably contributed to the significant depletion of TE insertions in coding regions, as such insertions, presumably deleterious, would lead to their elimination from the tissue by clonal competition. Accordingly, insertion site enrichment in genic regulatory regions (UTRs and introns) could suggest a beneficial impact of transposition on the positive selection of cells with somatic TE insertions. Enrichment of *de novo* insertions in open chromatin could, at least partially, be explained by physical DNA accessibility. However, we cannot exclude that other, yet unknown mechanisms also act at the pre-insertion level to direct retrotransposition away from exons and silent chromatin but towards non-coding genic regions and active chromatin.

### Consequences of somatic transposition

The impact of transposition on the biology of somatic tissues is under debate, as is its contribution to disease and aging (Faulkner and Garcia-Perez 2017; Chuong et al. 2017; Dubnau 2018). Here, we report evidence for somatic transposition with a functional impact on an adult tissue, by retrotransposon insertions into a tumor-suppressor gene *Notch.* As spontaneous neoplasia are isolated based on the characteristic *Notch* loss of function phenotype, and in those samples we found no other somatic events genome-wide that could explain inactivation of the Notch pathway (see also Riddiford et al, submitted), it is very likely that the somatic LTR retrotransposon insertions in *Notch* where indeed causative for neoplasia formation. Similarly, in mice somatic LTR element insertions causing oncogene or cytokine gene activation have been previously reported (Mietz et al. 1992; Howard et al. 2008). In human somatic *de novo* L1 retrotransposition activating oncogenic pathways has been documented in colorectal cancer (Miki et al. 1992; Scott et al. 2016) and hepatocellular carcinoma (Shukla et al. 2013), leading to a hypothesis that individual variation in somatically active elements could represent a novel form of cancer risk (Scott et al. 2016).

In addition to *Notch* inactivating events, many genome-wide retrotransposon insertions identified by us occurred in genic regions. We uncover exonic integrations, which would most likely lead to gene inactivation, as well as insertions in intronic or UTR sequences. Numerous examples of germline transposition show that TE insertions in non-coding genic regions can affect, both positively or negatively, target gene expression (Shen et al. 2011; Gong and Maquat 2011; Mateo et al. 2014; Ong-Abdullah et al. 2015; Ding et al. 2016; Van’t Hof et al. 2016). Among the genes hit by retrotransposition in our datasets, some are directly implicated in the tissue physiology, including regulators of ISC proliferation and differentiation. As for the *Notch* gene, in male flies, an X-chromosome insertion would result in a modification of a single available gene copy. In contrast, when two alleles are present (autosomes and female X chromosome), a somatic insertion would likely inactivate only one of them, limiting the functional impact. However, such insertions could still result in hypomorphic phenotypes. Moreover, unaffected alleles could be lost due to secondary genetic events of loss of heterozygosity, which we have shown to occur frequently in the fly gut (Siudeja et al. 2015; Siudeja and Bardin 2017). Finally, apart from directly affecting coding regions, non-genic transposable element insertions occurring in open chromatin could contribute cis-regulatory elements acting on neighboring or even distant genes leading to gain-of-function or mis-expression, as previously demonstrated for germline transposition (reviewed in (Chuong et al. 2017)). Hence, we hypothesize that transposition acting on genes or regulatory regions in the ISCs could influence stem cell fitness. By doing so, TE mobility could influence the clonal selection in the tissue by eliminating some and favoring other stem cell genomes.

Since somatic transposition may have functional consequences and contribute to diseases or aging, understanding how somatic transposition is controlled and how tissue-specificity arises, is of keen interest. We provide evidence that *Drosophila* will be an insightful model system for addressing mechanisms of somatic TE control and physiological consequences of somatic transposition.

## Materials and Methods

### Experimental techniques

#### Drosophila stocks and husbandry

*Pros>2XGFP* adults were obtained by crossing *w-*; *Pros*^V1^*GAL4/TM6BTbSb* females (J. de Navascués) with *w-*; *UAS-2XGFP;* males (Bloomington). *Dl>nlsGFP* were obtained by crossing *w-*; *DlGal4/TM6TbHu* (Wang et al. 2014) females with *w-*; *UAS-nlsGFP* males (Bloomington). Flies were maintained on a standard medium at 25°C with a day/night light cycle. For crosses, 10–15 females were mixed with males in standard vials. Progeny were collected over 2–4 days after eclosion. Adults were aged in plastic cages (mixed males and females) (10 cm diameter, 942 ml, 700–900 flies/cage) with freshly yeasted food provided in petri dishes every 2–3 days. Every 7 days, flies were transferred to clean cages.

#### Tissue isolation and short-read DNA sequencing

6-7-weeks-old *Pros>2XGFP* or *Dl>nlsGFP* males were used to isolate spontaneous neoplasia from unfixed tissues. Clones where identified by accumulation of GFP-positive cells. The midgut region containing an estimated 30%–80% neoplastic cells was manually dissected and transferred immediately to a drop of the ATL Buffer (Qiagen) for subsequent DNA isolation. Neighboring control gut tissue as well as the fly head and the fly thorax were also dissected. Genomic DNA was isolated with the QIAamp DNA MicroKit (Qiagen) according to the manufacturer’s protocol for processing laser-microdissected tissues. DNA quantity was measured with Qubit dsDNA High Sensitivity Assay Kit (Thermo Fisher Scientific). Genomic DNA libraries were prepared with the Nextera XT protocol (Illumina) using 0.6 ng of starting material. Whole-genome 2X100 bp or 2X150 bp paired-end sequencing was performed on HiSeq_2500 or Novaseq 6000 (Illumina). Supplemental Table S1 provides basic sequencing statistics for all samples used in this study.

#### PCR validation of somatic TE insertions in Notch and Sanger sequencing

We designed PCR primers down- and upstream of identified candidate insertions to amplify either a full-length TE insertion or a short wild-type genomic DNA fragment. PCR was performed on 0.5 – 1 ng of genomic DNA with the LongAmp Taq DNA Polymerase (New England Biolabs) using standard PCR reaction mix and long extension times (9 minutes at 65°C).

The following primers were used to generate Fig 1D:

R43-rover: GCAGCATTTGGTCCAAACGTT and ATAAAATGCGCCACAAGACGAG
R9-rover-ex: CGCGCAAGGATAATTGGATGG and GCGTAGTCTTATGGCCTAGTG
R9-rover-int: CCAGGCTGCAATTACTTTAATT and TGTAAAATGCAAGCGGAATGC

For Fig 1E, the DNA fragment containing the TE insertion was gel-purified and used as a template for a second PCR amplification using the same conditions as above. The amplified product was purified and Sanger sequencing was performed by Eurofins Genomics, using the following primers: GCAGCATTTGGTCCAAACGTT and CGTAGAGATTAGAGAATTACA.

#### RNA and small RNA isolation and sequencing

For RNA isolation, gut and head tissues from 1-week-old flies were dissected in cold, RNase-free PBS, transferred to 100L of TRIzol Reagent (Thermo Fisher Scientific), homogenized with a plastic pestle and snap frozen in liquid nitrogen for storage at −80°C. Upon thawing, samples were further processed according to the TRIzol Reagent manufacturer’s protocol. Purified RNA was treated with DNase (Ambion) for 1hr at 37°C, further purified with phenol-chloroform extraction and isopropanol precipitation, and resuspended in RNase-free water. All samples had A260/280 ratios above 1.9 and A260/230 ratios above 2.0. RNA integrity was checked on Bioanalyzer (Agilent) using the Agilent RNA 6000 Nano Kit and concentrations were assayed with the Qubit RNA Broad Range Assay Kit (Thermo Fisher Scientific). For the transcriptome analysis, 700ng of total RNA was used to prepare libraries according to the TruSeq Stranded mRNA protocol (Illumina). Samples were processed in biological triplicates. 2X100 bp paired-end sequencing was performed on Novaseq (Illumina). Small RNA fractionation and sequencing was performed by Fasteris, SA (Geneva, CHE). Briefly, after 2S rRNA depletion and PAGE gel-sizing for 18-30-nt fragment size, libraries were prepared according to the TruSeq small RNA Kit (Illumina) and sequenced on NextSeq 500, in 1×50bp single-end mode. Samples were processed in biological duplicates.

#### High molecular weight genomic DNA isolation and long-read sequencing

For sequencing of non-neoplastic tissues, we isolated guts from 25-days-old female flies without any visible neoplasia along with the fly heads. Tissues were dissected in ice-cold, nuclease free PBS and snap-frozen in liquid nitrogen before DNA isolation. High molecular weight genomic DNA was isolated from pools of 60 guts or 60 heads with the MagAttract HMW DNA Kit (Qiagen) according to manufacturer instructions. gDNA was eluted with nuclease-free water. DNA integrity was verified on a 0.6% agarose gel and concentrations were measured with Qubit dsDNA Broad Range Assay Kit. All samples had A260/280 ratios above 1.8 and A260/230 ratios above 2.0. Libraries were prepared with 800ng of DNA following the 1D Genomic DNA by Ligation Protocol (SQK-LSK109, Oxford Nanopore Technologies). Sequencing was performed on MinION using R9.4.1 flow cells (Oxford Nanopore Technologies) and 48hr-long sequencing runs. Supplemental Table S1 provides basic sequencing statistics for all samples used in this study.

### Computational analysis

#### Putative TE insertion detection from short-read DNA sequencing

The following applies to all Illumina short read paired end DNA sequencing datasets. Adapter sequences were trimmed using fastp version 0.19.5 (Chen et al. 2018). Trimmed reads were aligned to release 6.13 of the *Drosophila Melanogaster* reference genome (FlyBase) using bwa-mem version 0.7.17. bwa-mem parameters were default parameters, except for -Y (use soft clipping for supplementary alignments) and -q (don’t modify mapQ of supplementary alignments). Trimmed reads were also aligned to *Drosophila melanogaster* TE family consensus sequences (https://github.com/bergmanlab/transposons) using bowtie2 (Langmead et al. 2009) version 2.3.4.3. Duplicate reads were marked using picard markdup 2.18.2 (http://broadinstitute.github.io/picard/). Genome alignments and TE alignments are inputs to the readtagger command of the readtagger package (https://github.com/bardin-lab/readtagger). readtagger writes SAM tags for alignments where either the alignment or the mate of the alignment also aligns to a TE or other non-reference genome sequence. Tags contain information about the alignment (TE reference, alignment start, alignment end, query start, query end, and alignment orientation), can be visualized in IGV and are used to locate potential non-reference TE insertions. Next, the findcluster command takes the tagged alignment files and iteratively splits and groups tagged reads within a distance that corresponds to the 95% interval of the insert distance into clusters based on their alignment orientation and clipped sequences, and annotates if any cluster shows signs of a target site duplication (TSD). These unfiltered clusters are further linked to soft-clipped sequences at the 5’ and 3’ ends of putative insertions, so that the presence of a particular clipped sequence at a given genomic position can be used as a proxy for determining whether reads are congruent with an insertion or not.

The output of the findcluster step is a GFF file containing putative insertions and soft-clipped sequences and their genomic position as well as a BAM file containing only aligned reads assigned to a cluster (this includes alignments that support an insertion and alignments that support the reference).

#### Filtering putative TE insertions to obtain non-reference somatic TE insertions

findcluster outputs for each sample were filtered to retain only insertions that contain both mate and split read support. These putative insertions were then processed with the confirm_insertions command of the readtagger package. This command takes as input a file containing pre-filtered putative insertions in GFF format, a set of all putative insertions from all samples and a set of insertions from all samples that can be considered a panel of normal. To detect somatic TE insertions for a particular tumor dataset the panel of normals are all head datasets. Inversely, to detect somatic head insertions, the panel of normal are all tumor datasets. confirm_insertions links insertions from all samples using overlapping clipped sequences, genomic location and the family of a putative TE insertion. Putative somatic TEs were those insertions that were not found within the panel of normal. We further required a valid target site duplication to be present. For each candidate somatic TE insertion we generated an IGV screenshot that includes 500 nucleotides up- and downstream of each insertion. Upon visual inspection of screenshots putative insertions that were likely incorrectly called due to imprecise annotation of either the putative insertion or the control insertion were discarded. The lists of all identified putative somatic insertions can be found in Supplemental Tables S4 (gut clonal samples) and S5 (head samples).

#### Filtering putative TE insertions to obtain non-reference germline insertions

To estimate the rate of germline transposition, datasets were analyzed as for somatic TE insertions, but we treated each family individually, where the panel of normal constituted all other families. The retained insertions were then private to the family being analysed.

#### Comparison of somatic TE allele frequencies to neoplasia-initiating events

Allele frequencies of somatic insertions were estimated from the total pool of read pairs that overlap a putative insertion. Allele frequencies of neoplasia-initiating events were taken from a companion paper addressing structural variation in the same model system (Riddiford, et al, submited). The exception to this were samples P15, P47 and D5, where somatic TE insertions in *Notch* were assigned as neoplasia initiating events. For sample P15, with two somatic insertion in *Notch*, the insertion with a higher allele frequency was set as the putative *Notch* inactivating event. We plotted the estimated allele frequency of insertions as dots and the tumor initiating event as a bar. We adjusted the allele frequency of TE insertions on autosomes by multiplying with a factor of 2.

#### Long-read sequencing data analysis

Nanopore reads were basecalled using guppy version 3.2.4. Since read length and sequencing depth is not uniform for long-read datasets and this can affect the number of full-length TE insertions that are detectable, reads were normalized using the normalize_readsizes of the readtagger package. All analysed nanopore libraries therefore have the same read-size distribution and sequencing depth. Reads were aligned to release 6.13 of the *Drosophila melanogaster* reference genome using minimap2 version 2.17 (Li 2018) with the -Hk19 preset for Nanopore reads and the -Y flag. Alignments with a mapping quality below 40 were discarded with samtools view. extract_variants from the readtagger package was used to check all soft-clipped or insert sequences for homology to TEs using mappy. Aligned positions around soft-clipped or insert sequences were written to a new alignment file along with a tag that describes the TE alignment. For soft-clipped sequences, a single aligned N-nucleotide was written out together with the soft-clipped sequence. For inserts, the insert sequence was written out using 1 flanking N nucleotide at each site. Alignment files were then parsed into a tabular format and analysed to find unique full-length transposable element insertions using ipython notebooks available at https://github.com/bardin-lab/somatic-transposition-fly-intestine. The lists of all identified singleton insertions can be found in Supplemental Tables S6 (pooled gut samples) and S7 (pooled head samples).

#### Motif analysis at integration sites

Motifs were determined by extracting 10 flanking nucleotides upstream and downstream of each insertion using bedtools slopbed and bedtools getfastabed (Quinlan and Hall 2010) and running meme version 5.0.5 (Bailey et al. 2009) on the resulting multi-fasta file. meme parameters were -dna for DNA alphabet, -revcomp which checks the reverse complement for motifs, -pal for checking for palindromes and a motif width between 9 and 50 (−minw 8, maxw 50).

#### Genome features enrichment analysis

Pre-analyzed modencode datasets (The modENCODE Consortium et al. 2010) were downloaded and lifted over to release 6.13 of the *Drosophila* genome. DamID peaks (Gervais et al. 2019) were downloaded from GSE128941. Overlap was analyzed in ipython notebooks available at https://github.com/bardin-lab/somatic-transposition-fly-intestine. pybedtools fisher (Dale et al. 2011) was used to determine enrichment and significance of overlap. Correlations with p-value<0.0001 and −1.5>enrichment>1.5 were considered significant. The overlap between genes and TE insertion sites (Fig 6C) was calculated with bedtools windowbed using +/− 500 bp window size.

#### RNAseq data analysis

For RNAseq analysis reads were trimmed off their adapters using Atropos (Didion et al. 2017) and quasi-mapped against the *Drosophila* reference transcriptome (release 6.13, Flybase) supplemented with family-level TE sequences (https://github.com/bergmanlab/transposons) using Salmon version 0.14.1 (Patro et al. 2017) and RPM values were reported in Fig 4. Differential expression analysis (Fig S2) was performed on previously published datasets (Patel et al. 2015) using DESeq2 (Love et al. 2014). TE transcripts with an adjusted p-value < 0.01 were considered differentially expressed. We used the plot_coverage command of the readtagger package to create coverage plots.

#### Small RNA analysis

Sequencing adapters were trimmed using fastp version 0.19.5. Reads were aligned to family level TE sequences using HISAT2 (Kim et al. 2019) version 2.1.0. Size distributions were plotted using ipython notebooks. Ping-pong signatures were calculated using the Small RNA Signatures tool, version 3.1.0 (Antoniewski 2014).

## Data availability

Datasets used for this study are available under the following accession numbers:

ArrayExpress, E-MTAB-3917 – whole-genome neoplasia/head control sequencing (samples P1-P6; (Siudeja et al. 2015))
NCBI, Bioproject, PRJNA641572 – whole genome neoplasia/head control sequencing (samples P7-P66 and D1-D8; this study)
XXXXXXXX – Nanopore ONT sequencing of pooled guts/heads, RNAseq of pooled guts and small RNAseq of pooled guts and ovaries (this study)

## Acknowledgements

High-throughput sequencing has been performed by the ICGex NGS platform of the Institut Curie supported by the grants ANR-10-EQPX-03 (Equipex) and ANR-10-INBS-09-08 (France Génomique Consortium) from the Agence Nationale de la Recherche (“Investissements d’Avenir” program), by the Canceropole Ile-de-France and by the SiRIC-Curie program - SiRIC Grant « INCa-DGOS-4654”. We thank C. Antoniewski, P. Hollyoak, T. Hall and W. Hamitou for their contribution to some experiments and data analysis; N. Da Cruz and B. Leveille Nizerolle for technical assistance; as well as members of the Bardin team, N. Servant, J. Waterfall, D. Bourc’his, M. Greenberg and P.-A. Defossez for discussions and comments on the manuscript. This work was supported by grants from Fondation pour la Recherche Médicale (A.B., DEQ20160334928), as well as funding from the program “Investissements d’Avenir” launched by the French Government and implemented by ANR, ANR SoMuSeq-STEM (A.B), Labex DEEP (ANR-11-LBX-0044), IDEX PSL (ANR-10-IDEX-0001-02 PSL) and ICGex STEM-SOM-GERM (A.B. and K.S). Salary support of K.S. is from Inserm; A.J.B by CNRS; N.R is through a grant from Fondation ARC, and B.B by ENS Lyon.

## Author Contributions

K.S., M.v.d.B. and A.J.B. designed the study and analyzed the data. M.v.d.B. developed software and performed all bioinformatic analysis. K.S. performed a majority of the experiments. B.B. collected samples for the Delta>nlsGFP background; M.S. obtained data for Supplemental Fig S4; and A.W. contributed to the long-read sequencing experiments. N.R. provided variant frequencies of neoplasia initiating events for Fig 3B. S.L. prepared samples for sequencing. K.S. and A.J.B. wrote the manuscript with contribution from M.v.d.B. and other authors.

The authors declare no competing interest.

