## Supplemental Figures for "Somatic transposition in the *Drosophila* intestine occurs in active chromatin and is associated with tumor suppressor gene inactivation"

* Equal contribution

**Supplemental Figures:**

**Supplemental Figure S1.** Rare somatic transposition events are found in head samples

**Supplemental Figure S2.** Depletion of Notch in gut progenitor cells has a minor effect on TE expression levels

**Supplemental Figure S3.** The analysis of small RNA fractions isolated from ovaries

**Supplemental Figure S4.** The analysis of germline TE insertions

**Supplemental Figure S5.** Enrichments of putative somatic singleton insertions identified with the long-read sequencing of bulk gut DNA

**Supplemental Figure S6.** Correlations of somatic insertion sites from the short-read sequencing of clonal samples with modENCODE tracks for *Drosophila* larvae.

**Supplemental Tables:**

**Supplemental Table S1.** All whole-genome short- and long-read DNA samples used for this study.

**Supplemental Table S2.** List of genes with somatic TE insertions recovered from the *Pros>2xGFP* clonal gut samples (Illumina sequencing)

**Supplemental Table S3.** List of genes with singleton insertions recovered from pooled *Pros>2xGFP* normal midguts (Nanopore ONT sequencing)

**Supplemental Table S4.** All putative somatic TE insertions (with a TSD) identified in the gut clonal samples with the Illumina sequencing.

**Supplemental Table S5.** All putative somatic TE insertions (with a TSD) identified in the head samples with the Illumina sequencing.

**Supplemental Table S6.** All singleton TE insertions (with a valid TSD) identified in the Pros>2xGFP gut pooled samples with the ONT Nanopore sequencing

**Supplemental Table S7.** All singleton TE insertions (with a valid TSD) identified in the Pros>2xGFP head pooled samples with the ONT Nanopore sequencing

Siudeja_Fig.S1

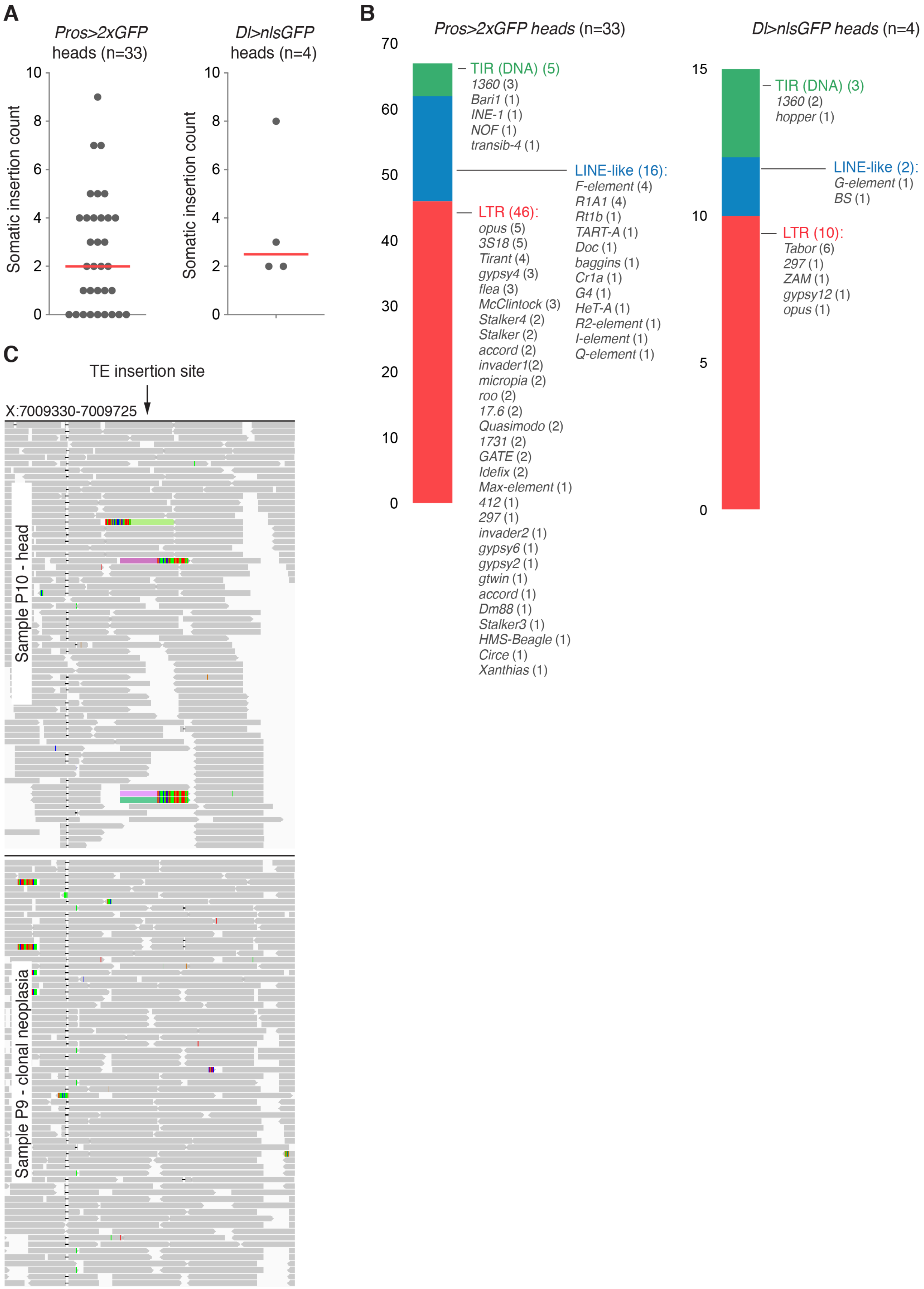


**Figure S1. Rare somatic transposition events are found in head samples**

**(A)** The frequency of head specific somatic insertion sites in the *Pros>2XGFP* and *Delta>nlsGFP* genetic backgrounds. Red lines represent median values. **(B)** The distribution of TE classes active in the heads of two genetic backgrounds studied. **(C)** The IGV screenshot of a head-specific TE insertion site from sample P10 (head) and it's neoplasia control (sample P9). Bars represent sequencing reads. The reads supporting the TE insertion are colored.

### Siudeja_Fig.S2

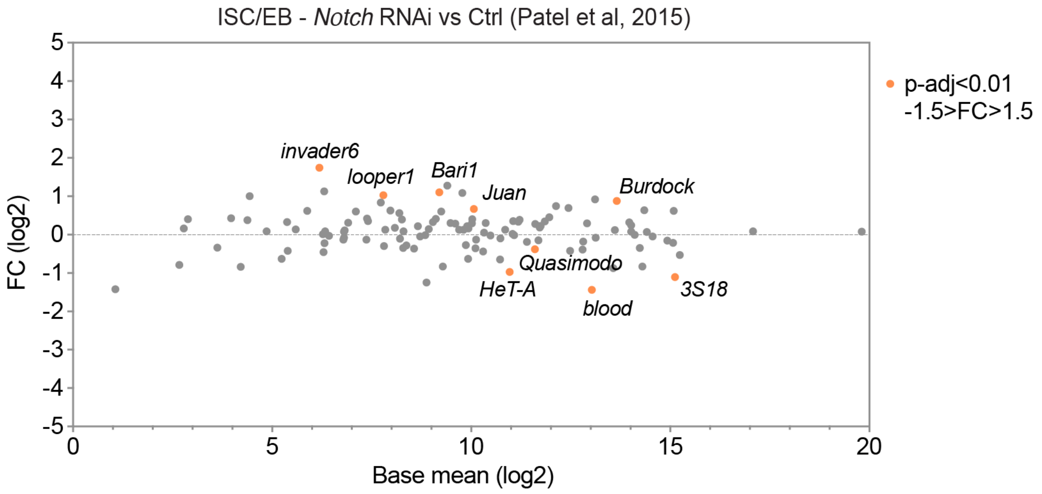


**Figure S2. Depletion of Notch in gut progenitor cells has a minor effect on TE expression levels.** Differential expression of TEs in FACS-sorted wild-type or *Notch*-depleted gut progenitor cells (*escargot* positive ISCs and enteroblasts). The data is from Patel et al, Nat Cell Biol, 2015. TEs with p(adj)<0.01 and -1.5>FC>1.5 were considered differentially expressed (labeled and marked in orange).

Siudeja_Fig.S3

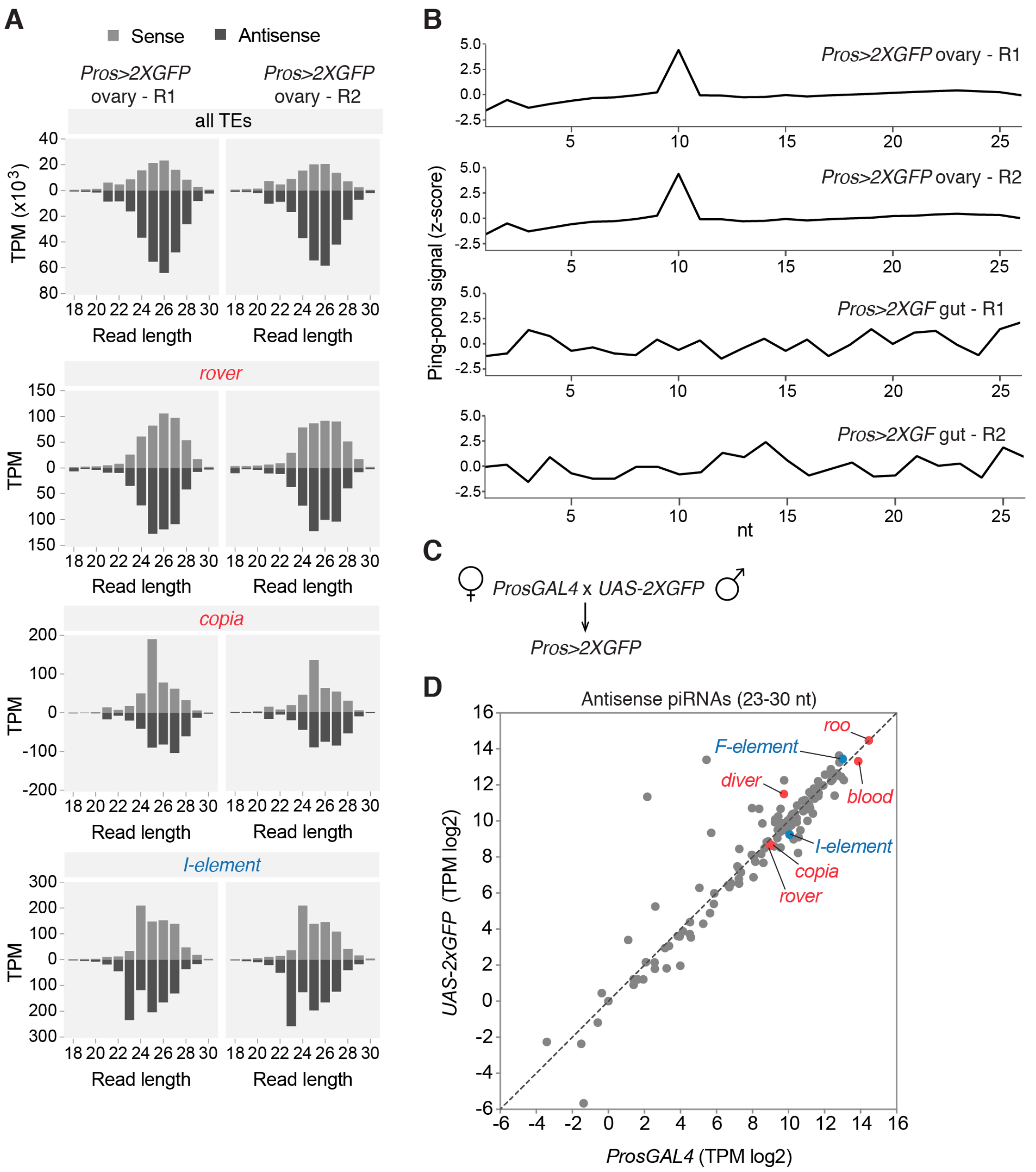


**Figure S3. The analysis of small RNA fractions isolated from ovaries**

**(A)** The size distribution of sense and antisense reads from *Pros>2XGFP* ovary small RNA fractions mapping to all TEs (upper panel) or selected TE families mobilizing in the gut. R1 and R2 are two biological replicates. **(B)** The complementary sense and antisense read overlap (z-score) calculated on the 23-30nt long small RNA populations from ovary and gut samples. The 10nt overlap detected in ovary, but not gut samples, is a signature of the piRNA "ping-pong" cycle. **(C)** Parental fly crossing scheme used to obtain the *Pros>2GFP* genotype used in this study. **(D)** Scatter plot showing normalized TE-mapping antisense piRNA levels from ovaries of two parental stocks used to obtain the *Pros>2xGFP* flies. TEs generating most of the somatic gut insertions are highlighted in red (LTR-elements) or blue (LINE-like).

**Siudeja_Fig.S4**

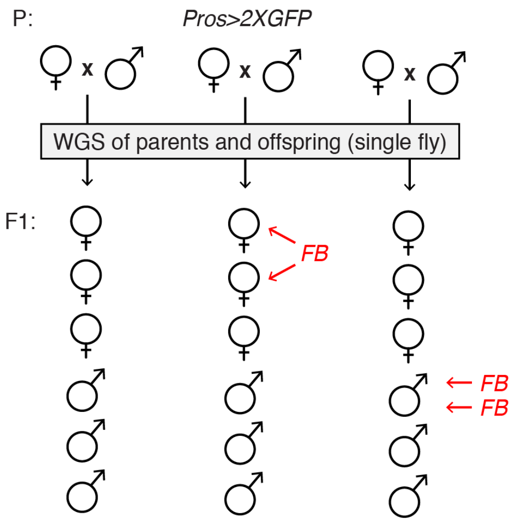


**Figure S4. The analysis of germline TE insertions**

Schematic representation of individual flies sequenced to detect germline TE insertions transmitted to the progeny (F1) of the *Pros>2xGFP* parents (P). Detected *de novo* germline TE insertions are indicated with horizontal red arrows. We detected 3 germline *de novo foldback* element insertions. Two of those were present in 1 male and one was detected in 2 sibling females. FB – *foldback* element.

**Siudeja_Fig.S5**

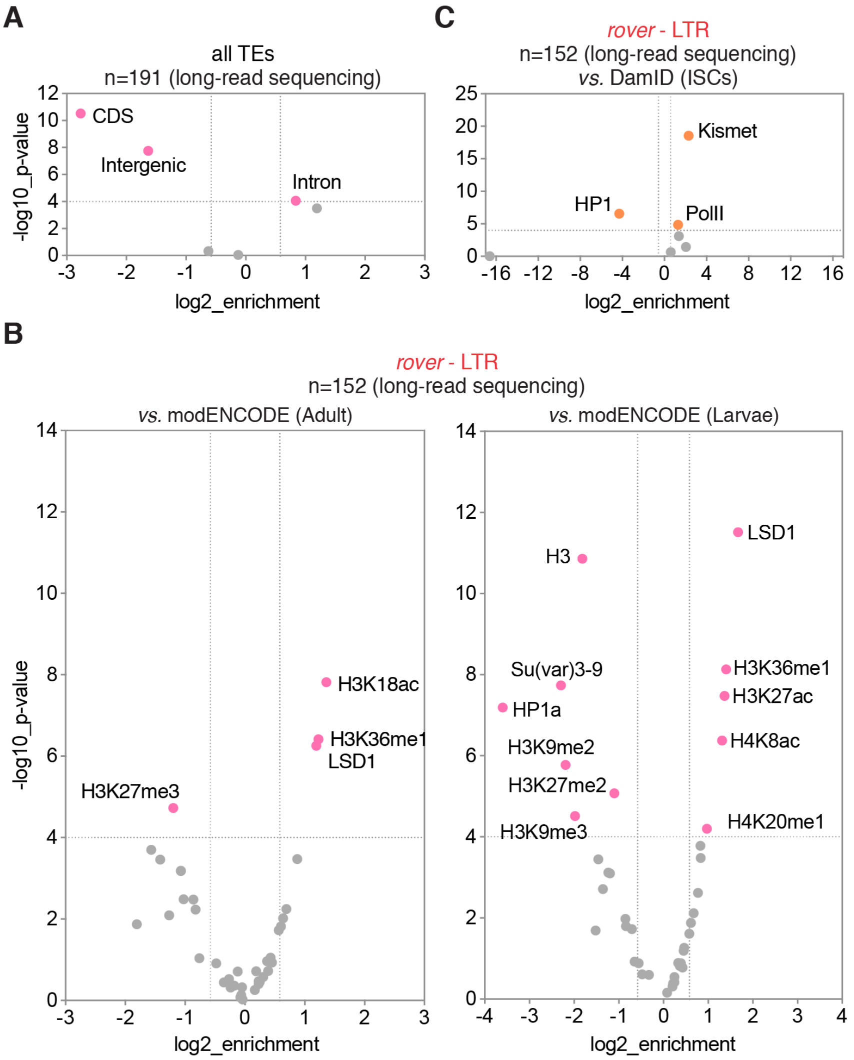


**Figure S5. Enrichments of putative somatic singleton insertions identified with the long-read sequencing of bulk gut DNA**

**(A)** Candidate singleton insertion sites were depleted from intergenic and exonic sequences and enriched in 3'UTR regions of the fly genome. **(B, C)** Correlations of singleton insertion sites of *rover* elements with modENCODE tracks for adult fly (**B**) and larval (**C**) tissues. **(D)** Correlations of singleton insertion sites of *rover* elements with DamID tracks for adult fly intestinal stem cells (ISC). Colored data points and labels highlight significant positive or negative correlations (p<0.0001, -1.5>enrichment>1.5).

### Siudeja_Fig.S6

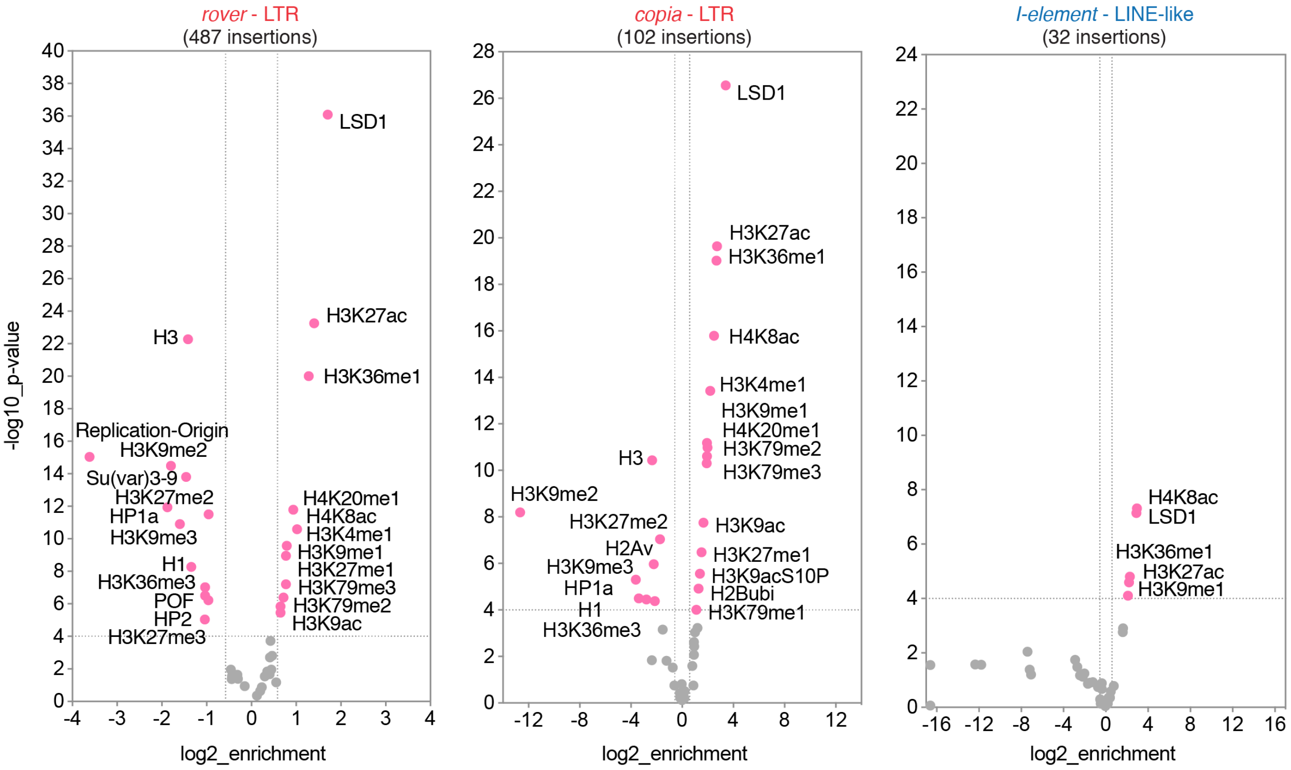


**Figure S6. Correlations of somatic insertion sites from the short-read sequencing of clonal samples with modENCODE tracks for *Drosophila* larvae.**

Three most represented TE families (*rover*, *copia* and *I-element*) are plotted. Colored data points and labels highlight significant positive or negative correlations (p<0.0001, -1.5>enrichment>1.5). Insertions from the *Pros>2XGFP* clonal gut short-read sequencing samples were used for all plots.
